# Modeling the dynamics of Let-7-coupled gene regulatory networks linking cell proliferation to malignant transformation

**DOI:** 10.1101/519165

**Authors:** Claude Gérard, Frédéric Lemaigre, Didier Gonze

## Abstract

The microRNA Let-7 controls the expression of proteins that belong to two distinct gene regulatory networks, namely a cyclin-dependent kinases (Cdks) network driving the cell cycle and a cell transformation network which can undergo an epigenetic switch between a non-transformed and a malignant transformed cell state.

Using mathematical modeling and transcriptomic data analysis, we here investigate how Let-7 controls the cdk-dependent cell cycle network, and how it couples the latter with the transformation network. We also determine whether the two networks can be combined into a larger entity that impacts on cancer progression.

Our analysis shows that the switch from a quiescent to a cycling state depends on the relative levels of Let-7 and several cell cycle activators. Numerical simulations further indicate that the Let-7-coupled cell cycle and transformation networks control each other, and our model identifies key players for this mutual control. Transcriptomic data analysis from the The Cancer Genome Atlas (TCGA) suggest that the two networks are activated in cancer, in particular in gastrointestinal cancers, and that the activation levels vary significantly among patients affected with a same cancer type. Our mathematical model, when applied to a heterogeneous cell population, suggests that heterogeneity among tumors results from stochastic switches between a non-transformed cell state with low proliferative capability and a transformed cell state with high proliferative property. The model further predicts that Let-7 may reduce tumor heterogeneity by decreasing the occurrence of stochastic switches towards a transformed, proliferative cell state.

In conclusion, we identified the key components responsible for the qualitative dynamics of two GRNs interconnected by Let-7. The two GRNs are heterogeneously involved in several cancers, thereby stressing the need to consider patient’s specific GRN characteristics to optimize therapeutic strategies.

## Introduction

Let-7 microRNAs control two distinct gene regulatory networks (GRNs) that regulate cell cycling and malignant transformation of breast cancer cells (Johnson et al., 2007;Iliopoulos et al., 2009). A cyclin-dependent kinases (Cdk) network controls the correct progression of the cell cycle along the G1, S, G2 and M phases (Morgan, 2007). Growth factors (GF) and E2F stimulate, while Let-7 down-regulates the expression of several components of this cdk-dependent cell cycle network (Bueno and Malumbres, 2011). Mathematical models focusing on post-translational regulations of cyclin/Cdk complexes were proposed to account for the dynamics of the Cdk network in mammals (Novak and Tyson, 2004;Gerard and Goldbeter, 2009). However, to our knowledge, no model has been proposed to study the impact of miRNAs on this network. Let-7 is also a key component of a GRN which promotes cell transformation in response to an inflammatory stimulus (Iliopoulos et al., 2009). This GRN is characterized by a positive feedback loop (PFL), where a transient inflammatory stimulus is sufficient to induce the cells to undergo a PFL-dependent epigenetic switch from a non-transformed state toward a permanently malignant transformed state. We previously proposed a model describing the dynamics of this transformation GRN (Gerard et al., 2014). Our model suggested that a transient inflammatory signal activates an irreversible bistable switch in the expression of the GRN components, eventually leading to a stable epigenetic switch allowing cells to display increased motility and invasiveness. In this GRN, Let-7 prevents cell transformation by inhibiting the translation of Interleukin-6 (IL6) and Ras, two drivers of the inflammatory feedback loop.

Let-7 being a component common to the cell cycle and transformation networks we now raise the following questions: how does Let-7 control the cdk-dependent cell cycle network; does Let-7 play a coupling role between the cell cycle GRN and the transformation GRN, and can the two GRNs be combined into a larger network that impacts on cancer progression? We address these issues using experiment-based mathematical modeling of the GRNs and by analyzing transcriptomic data from The Cancer Genome Atlas (TCGA). Our approach identifies the dynamics of the two GRNs and indicates that their activation is cancer-specific and heterogeneous from patient to patient, stressing the need to consider patient’s specific characteristics.

## Results

### Structure of the cell cycling and transformation networks

The structure of the Cdk network gives rise to a transient and sequential activation of the various cyclin/Cdk complexes, allowing for a correct progression through the different cell cycle phases (Fig. 1A and (Morgan, 2007;Gerard and Goldbeter, 2009)). The activity of cyclin D/Cdk4-6 ensures the transition G0/G1 and the progression in G1. Cyclin E/Cdk2 promotes the G1/S transition, while cyclin A/Cdk2 elicits progression in S and G2. Finally, cyclin B/Cdk1 brings about G2/M transition and the entry of cell into mitosis (Fig. 1A). In the model, Let-7 represses each cyclin/Cdk complex.

**Figure 1.**
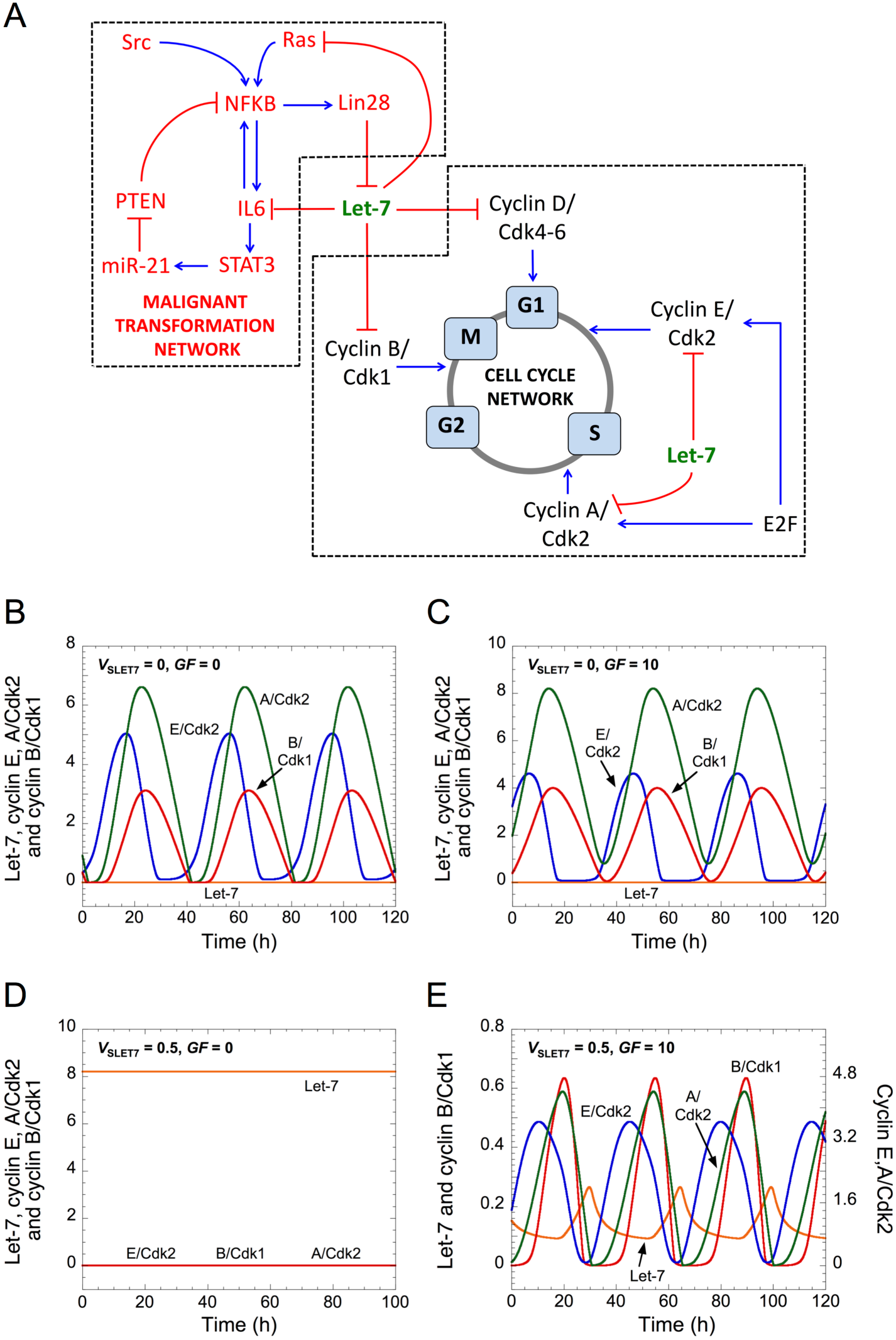
Let-7 and GF control cell cycle progression. (A) Scheme of both GRNs coupled by Let-7. (B-E) Temporal evolution of Let-7, cyclin E/Cdk2, cyclin A/Cdk2 and cyclin B/Cdk1 are shown in the absence (*V*_SLET7_ = 0 in B, C) or in the presence of Let-7 (*V*_SLET7_ = 0.5 in D, E), and in the absence (*GF* = 0 in B, D) or in the presence of GF (*GF* = 10 in C, E). Parameter values are as in Supplementary Table 3. A detailed scheme of both GRNs that includes all regulations considered in the model is in Supplementary Fig. 1.

Let-7 is also at the core of a positive feedback loop (PFL) in a malignant transformation network (Iliopoulos et al., 2009). Indeed, a transient inflammatory signal mediated by the oncoprotein Src stably activates NF-κB, which promotes Lin28, IL6 and STAT3 activation (Iliopoulos et al., 2010;Fofaria and Srivastava, 2015).

Thus, we propose that Let-7 couples gene regulatory networks linking cell proliferation to malignant transformation.

### Relative levels of Let-7 and growth factors control cell cycling

To analyze the impact of Let-7 on the Cdk network dynamics, we built a qualitative mathematical model of cell cycle regulation by Let-7 (see model’s structure in Figs. 1A and in Supplementary Fig. 1 for a detailed description highlighting all regulatory interactions included in the model). The model is an extension of an earlier model of the Cdk network that accounted for the dynamics of the mammalian cell cycle (Gerard and Goldbeter, 2011). It now explicitly considers the mRNA forms of each cyclin, enabling us to incorporate Let-7-mediated post-transcriptional regulations of cyclin synthesis. Let-7 represses the synthesis of multiple activators of the cell cycle, such as cyclins D and A and Cdk2/4/6 (Bueno and Malumbres, 2011). For sake of simplicity we consider that Let-7 directly represses the translation of cyclins D, E, A and B, by forming an inactive complex with their respective mRNA. In addition, GF promote the synthesis of cyclin D, eliciting the G0/G1 transition and the entry of the cell into the cell cycle, while E2F activates synthesis of cyclins E and A.

The model of the cell cycle network is composed of a set of kinetic equations describing the temporal evolution of the levels of each component of the network. It includes the mRNAs of cyclin D, E, A, and B; the active form of E2F; the various cyclin/Cdk complexes: cyclin D/Cdk4-6, cyclin E/Cdk2, cyclin A/Cdk2 and cyclin B/Cdk1; and the active form of the Anaphase-Promoting Complex, APC, which triggers degradation of cyclins A and B at the end of mitosis (Supplementary Fig. 1). Variables are defined in Supplementary Table 1, the kinetic equations are in Supplementary Table 2, and the parameters are defined in Supplementary Table 3.

As a consequence of its regulatory structure the network self-organizes with sustained oscillations in the activity of the various cyclin/Cdk complexes, which correspond to successive rounds of cell cycling (Fig. 1B-E). In the absence of both Let-7 and GF, cells proliferate and sustained oscillations of the various cyclin/Cdk complexes develop (Fig. 1B). This situation bears similarity with transformed or cancer cells, which are often characterized by downregulation of Let-7 and signal-independent growth (Sotiropoulou et al., 2009;Hanahan and Weinberg, 2011). Starting from that condition, an increase in GF maintains cell proliferation (Fig. 1C), while an increase in Let-7 suppresses cell proliferation (Johnson et al., 2007). The latter case is characterized by a stable steady state, with low levels of each cyclin/Cdk (Fig. 1D). Finally, starting from that steady state, an increase in GF permits to recover cell proliferation (Fig. 1E).

The respective impact of Let-7 and GFs can be visualized in a two-parameter plane where the dynamical behavior of the cell cycle network is represented as a function of the synthesis rate of Let-7, *V*_SLET7_, and the level of growth factors, *GF* (Fig. 2A). For large values of *V*_SLET7_ (high levels of Let-7), the Cdk network tends to a stable steady state corresponding to cell cycle arrest regardless of GF levels. This corroborates experimental results showing that Let-7 represses cell proliferation (Johnson et al., 2007;Zhu et al., 2015). In contrast, for small values of *V*_SLET7_ (low levels of Let-7), the cell cycle network is characterized by sustained oscillations. The temporal evolution of cyclin E/Cdk2 and cyclin B/Cdk1 corresponding to conditions A to F in Fig. 2A are represented in Supplementary Fig. 2A-F, respectively. The sustained oscillations in cyclin E/Cdk2 with oscillations in cyclin B/Cdk1 of very small amplitude might correspond to endoreplication (Edgar et al., 2014) where multiple rounds of DNA replication occur without entry into mitosis (see temporal evolution in Supplementary Fig. 2D, which corresponds to condition D in Fig. 2A). Previous theoretical studies already showed that the regulatory structure of the cell cycle network in mammals is capable of generating endocycles (Gerard and Goldbeter, 2009).

**Figure 2.**
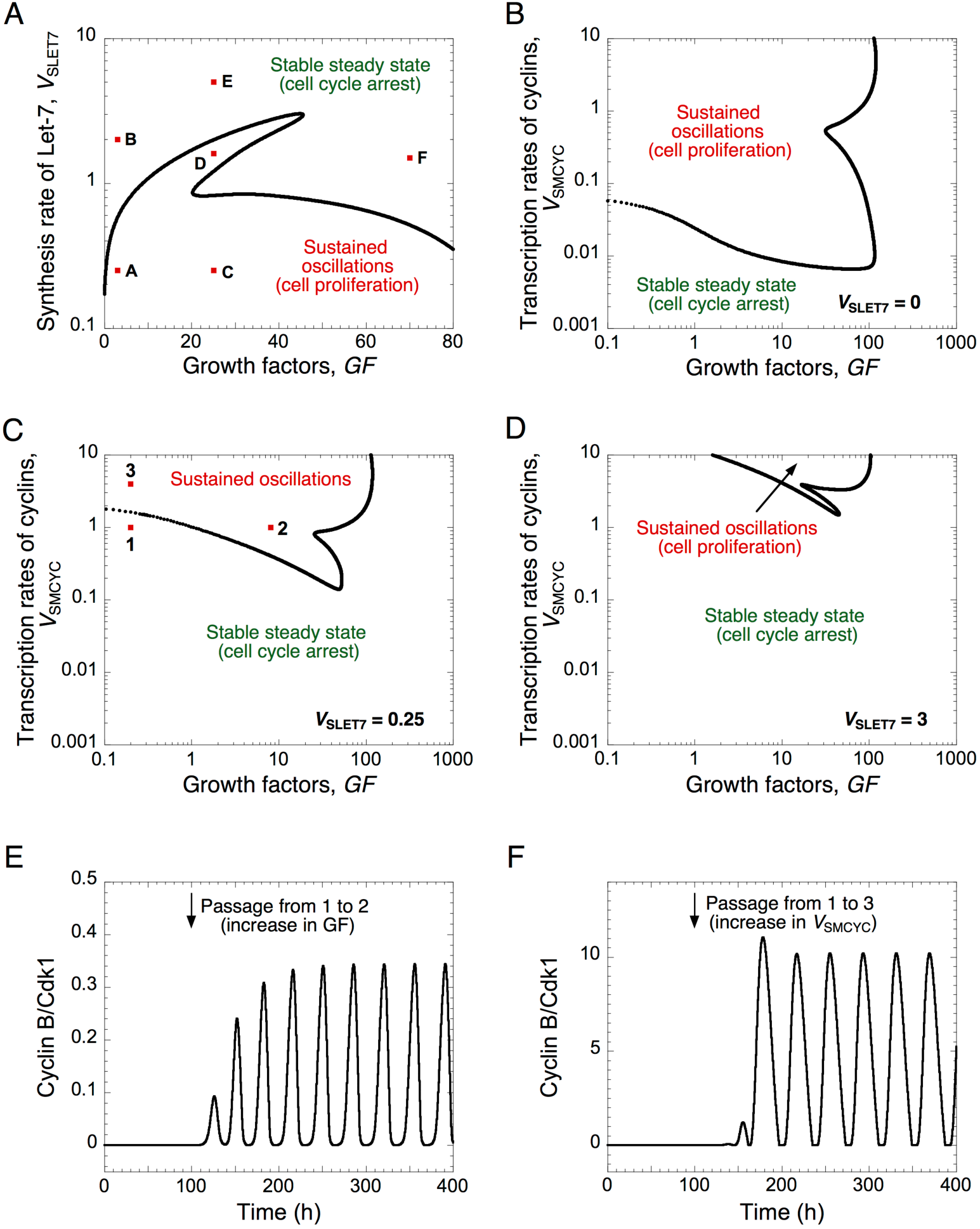
A balance between Let-7 and cell cycle activators determines the quiescence and proliferative state. (A) The Cdk network dynamics, i.e. sustained oscillations *versus* stable steady states, is represented in a two-parameter plane defined by the synthesis rate of Let-7, *V*_SLET7_, and the levels of GF. The Cdk network dynamics are indicated in a two-parameter plot as a function of the synthesis rate of the different cyclins, *V*_SMCYC_, and the levels of *GF* (B) in the absence of Let-7, *V*_SLET7_ = 0, (C) in the presence of intermediate, *V*_SLET7_ = 0.25, or (D) high Let-7 levels, *V*_SLET7_ = 3. (E, F) Temporal evolution of cyclin B/Cdk1 in the presence of GF overexpression (panel E where *GF* changes from 0.2 to 8, at t = 100h, which corresponds to the switch from conditions 1 to 2 in panel C), or in the presence of cyclin overexpression (panel F where *V*_SMCYC_ changes from 1 to 4, at t = 100h, which corresponds to the switch from 1 to 3 in panel C). Temporal evolution of cyclin E/Cdk2 and cyclin B/Cdk1 corresponding to conditions A to F in panel A are shown in Supplementary Fig. 2. Other parameter values are as in Supplementary Table 3.

The dynamics of the cell cycle network are further illustrated for different levels of Let-7 in a two-parameter plane defined by the synthesis rate of the cyclin/Cdk complexes, *V*_SMCYC_, and the level of GF, (Fig. 2B-D). By increasing Let-7, the domain of sustained oscillations, corresponding to cell proliferation, is reduced and limited to higher levels of cyclin/Cdk complexes (compare Fig. 2B, C, D where *V*_SLET7_ is equal to 0, 0.25 and 3, respectively). From a stable steady state, corresponding to quiescence (condition 1 in Fig. 2C), an increase in GF or an increase in the cyclin/Cdk levels may trigger the switch to sustained oscillations (see temporal evolution of cyclin B/Cdk1 in Fig. 2E and 2F).

We concluded that progression or arrest of the cell cycle is controlled by the relative levels of Let-7 and GF, or of Let-7 and the cyclin/Cdk complexes. Thus, when designing efficient anti-cancer strategies, the model stresses the importance to consider the relative, rather than the absolute, expression levels of network components displaying opposing effects on cell cycling.

### Let-7 couples the cell cycling and transformation networks

Let-7 belongs both to the cdk-dependent cell cycle GRN and the malignant transformation network (Fig. 1A). Therefore, we here determine the qualitative role of Let-7 as a coupling factor between the two GRNs, by analyzing the mutual impact of the GRNs on their respective dynamics.

Starting from a non-transformed, quiescent, cell state, defined by high Let-7 and low cyclin B/Cdk1 levels, a transient Src signal induced by inflammation triggers a stable down-regulation of Let-7, eliciting the switch of the cell cycle network from a stable steady state to sustained oscillations (Fig. 3A). Thus, transient inflammatory signals can promote persistent cell proliferation. On the opposite, as observed in the experiments (Iliopoulos et al., 2009), starting from a transformed and proliferative cell state, transient inhibition of Lin28, NF-κB or transient overexpression of PTEN stably impedes cell cycle progression (Fig. 3B-D).

**Figure 3.**
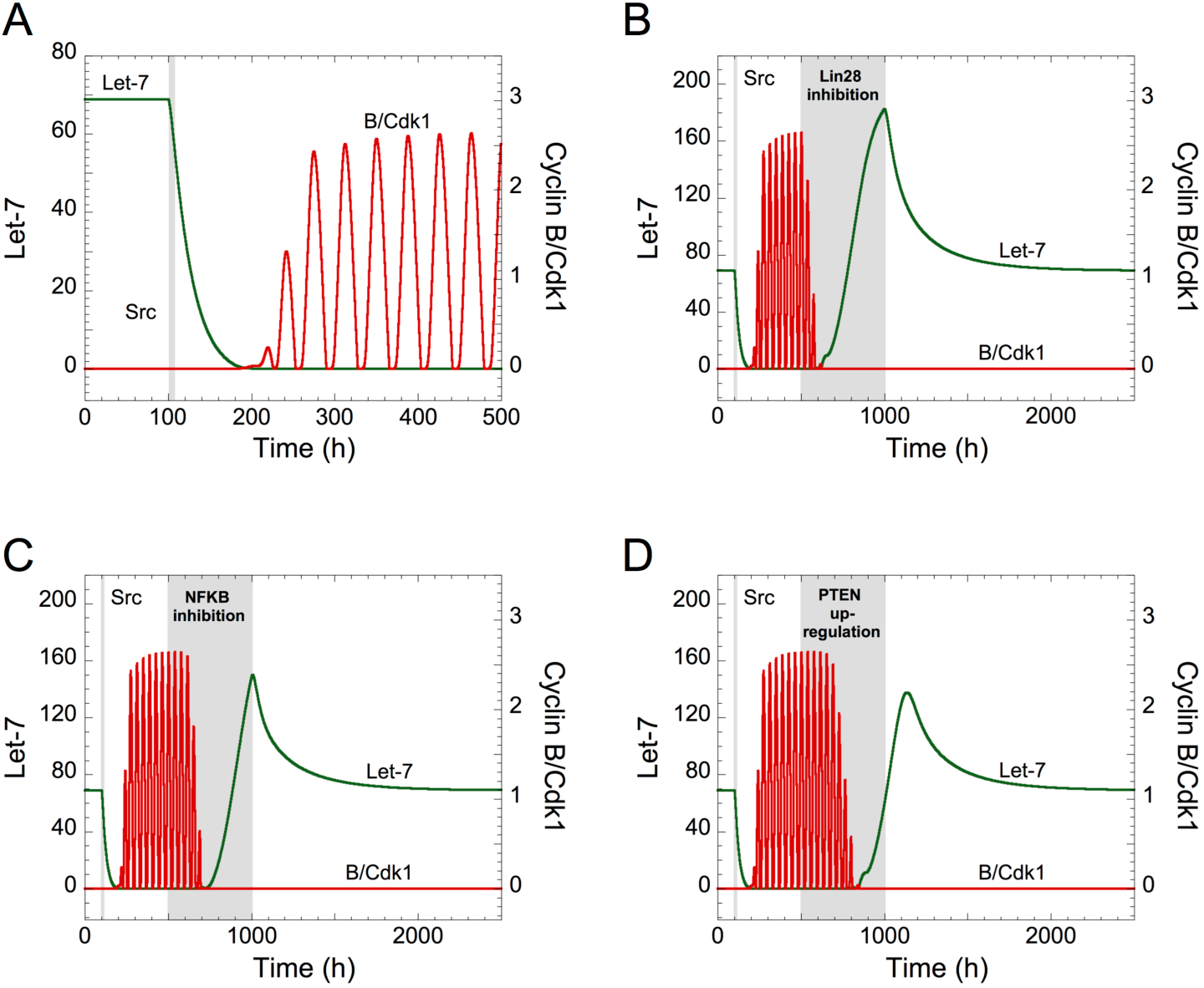
Let-7 is a central component between the proliferation and a malignant transformation network. (A) Effect of the inflammatory circuit on the Cdk network dynamics. (A-D) Starting from a non-transformed, quiescent, cell state, defined by high Let-7 and low cyclin B/Cdk1 levels, a transient increase in Src from t=100h to t=105h (grey area) triggers a stable epigenetic switch and cell proliferation characterized by Let-7 down-regulation and sustained oscillations of cyclin B/Cdk1. From that transformed, cell proliferative state, the temporal evolution of Let-7 and cyclin B/Cdk1 are illustrated in the presence of transient (B) Lin28 inhibition (*V*_SLIN28_ passes from 0.1 to 0 for 500h < t < 1000h), (C) NF-κB inhibition (*k*_AA1NFKB_ = *k*_AA2NFKB_ = *k*_AA3NFKB_ = 0 for 500h < t < 1000h), or (D) PTEN over-expression (*V*_SMPTEN_ passes from 0.001 to 10 for 500h < t < 1000h). Other parameter values are as in Supplementary Table 3.

We concluded that transient modifications in the expression of the components of the inflammation-dependent bistable transformation network can impact the long-term behavior of the cell cycling network when Let-7 couples the two GRNs.

To determine if the cell cycle network can modulate the dynamics of the transformation network, we simulated overexpression of all cyclins by increasing their synthesis rates, *V*_SMCYC_. Cyclin overexpression promotes uncontrolled cell proliferation of cancer cells (Gillett et al., 1994;Pok et al., 2013). Starting from a non-transformed cell state (high Let-7 levels), the model shows a stable down-regulation of Let-7 when *V*_SMCYC_ increases (Fig. 4A where *V*_SMCYC_ increases at t > 100h). The corresponding temporal evolution of Let-7 and cyclin B/Cdk1 is represented for different synthesis rates of cyclins (at t > 100h, *V*_SMCYC_ changes from 1.5 to 2.5 in Fig. 4B and from 1.5 to 12 in Fig. 4C). For weak overexpression of cyclins, the non-transformed quiescent cell state is maintained, while for stronger cyclin overexpression both GRNs are activated, i.e. downregulation of Let-7 and sustained oscillations of the cell cycle network (Fig. 4B-C).

**Figure 4.**
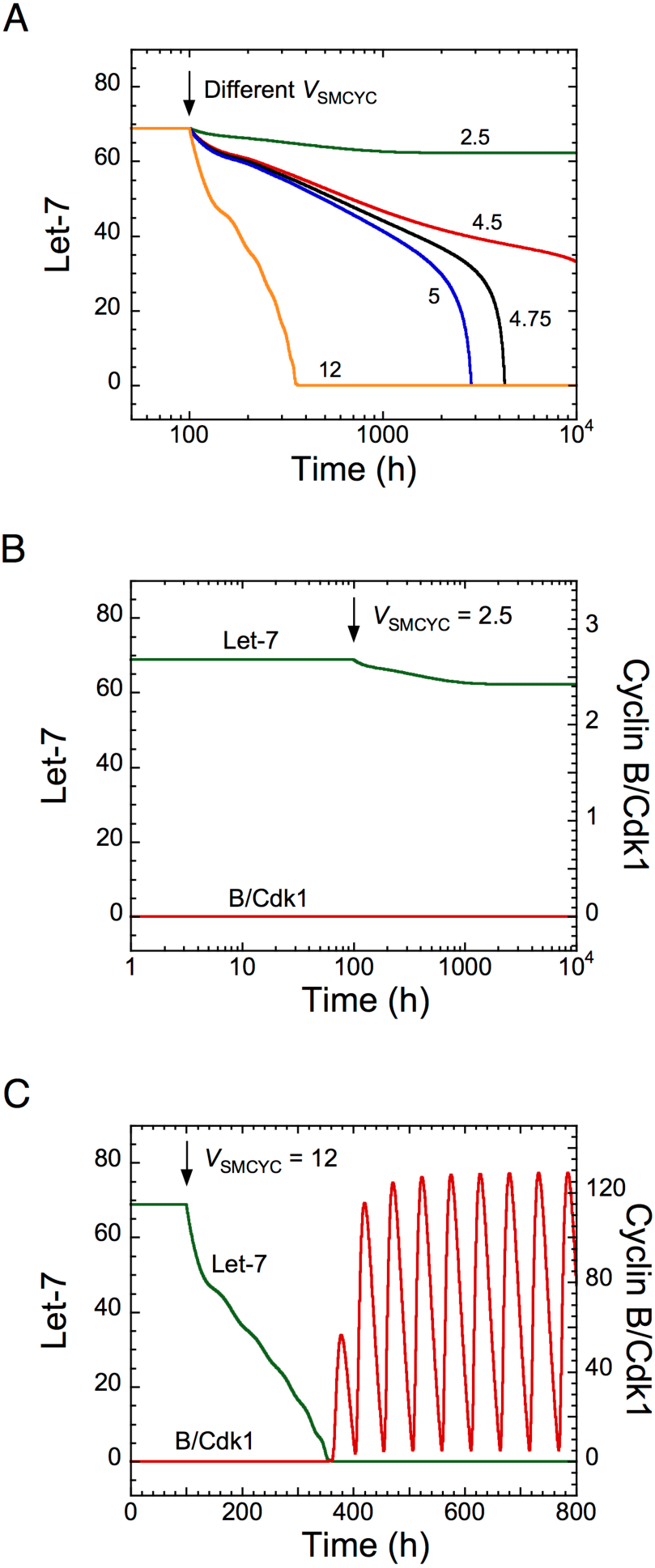
Overexpression of cyclins is predicted to impact on the dynamics of the transformation network. Temporal evolution of (A) Let-7, or (B, C) Let-7 and cyclin B/Cdk1 illustrated for different synthesis rates of cyclins, *V*_SMCYC_. From a non-transformed and quiescent cell state defined by high Let-7 levels and low levels of cyclin B/Cdk1, a sufficient increase in *V*_SMCYC_ (for t > 100h) down-regulates Let-7, allowing a stable switch to a transformed and cell proliferative state. Parameter values are as in Supplementary Table 3.

Temporal evolution of Let-7 and cyclin B/Cdk1 indicates that the switch to cell proliferation triggered by cyclin overexpression is irreversible because down-regulation of cyclins to their initial levels will not restore cell cycle arrest (Supplementary Fig. 3A, condition 1, 1200h < t < 1500h). This is the consequence of the irreversible bistable switch at the core of the transformation GRN (Gerard et al., 2014). The model predicts that a stronger decrease in cyclin synthesis can eventually stop cell proliferation, characterized by low, stable steady state levels of cyclin B/Cdk1 (Supplementary Fig. 3A, condition 3, t > 1500h). The corresponding temporal evolution of the expression levels of Lin28 and STAT3, two critical activators of the transformation GRN, is shown in Supplementary Fig. 3B. The model indicates that these cells (condition 3 for t > 1500h) might be invasive cells, defined by high levels of STAT3 and Lin28 and low Let-7 levels, which are in a quiescent state (low levels of cyclin/Cdk).

Moreover, the model predicts that a transient down-regulation of Lin28 can impede both cell proliferation and the transformation network (Supplementary Fig. 3C, condition 3, t > 1500h), where low, stable, levels of cyclin B/Cdk1 are present with high Let-7 levels. The corresponding temporal evolution of Lin28 and STAT3 expression levels is represented in Supplementary Fig. 3D. Here also, the modeling correlated well with the experimental observations showing downregulation of both cell proliferation and cell transformation after transient inhibition of Lin28 (Iliopoulos et al., 2009).

### Cancer type-specific activation of the proliferation and transformation networks

The coupling of the two networks raised the question of their potential combined involvement in cancer. To address this issue, we first defined a Cell Proliferation Index (*CPI*) and a Non-Transformed State Index (*NTSI*), where:

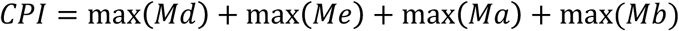

and

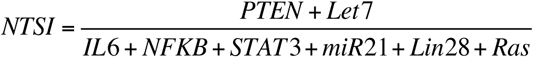

*CPI* is the sum of the maximal RNA levels of all cyclin/Cdk complexes, which is, as a first approximation, an indication of cell proliferation; *NTSI* is the ratio between the RNA expression levels of the inhibitors, i.e. Let-7 and PTEN, divided by the expression levels of the activators of the epigenetic transformation switch, i.e. IL6, NF-κB, STAT3, miR21, Lin28 and Ras. Such ratio characterizes the degree of cell transformation where a high value defines a non-transformed cell while a low value corresponds to transformed cells.

Since the components of the cell cycling and transformation networks are expressed in several tissues we verified if the networks are activated, i.e. if their components are consistently misexpressed in different types of cancer. We first calculated *CPI* and *NSTI* based on the RNA expression of the network components (Supplementary Table 5) from the non tumor (NT) and tumor samples (T) of TCGA cohorts of cholangiocarcinoma (CHOL: NT=9, T=36), stomach and esophageal carcinoma (STES: NT=46, T=600), hepatocellular carcinoma (LIHC: NT=50, T=369), stomach adenocarcinoma (STAD: NT=35, T=415), lung squamous cell carcinoma (LUSC: NT=51, T=501), bladder urothelial carcinoma (BLCA: NT=19, T=408), kidney renal clear cell carcinoma (KIRC: NT=72, T=534), breast carcinoma (BRCA: NT=112, T=1100), thyroid carcinoma (THCA: NT=57, T=510), kidney renal papillary cell carcinoma (KIRP: NT=32, T=290), kidney chromophobe (KICH: NT=25, T=66), and prostate adenocarcinoma (PRAD: NT=52, T=498) (Fig. 5A, B). Significant and consistently high *CPI* and low *NSTI* values, as compared to non-tumor conditions, were obtained in several cancer types, with largest variations in gastrointestinal cancers, i.e cholangiocarcinoma, hepatocellular and stomach and esophageal carcinoma. Interestingly, the proliferation and transformation networks do not seem to be activated in kidney and prostate adenocarcinoma.

**Figure 5.**
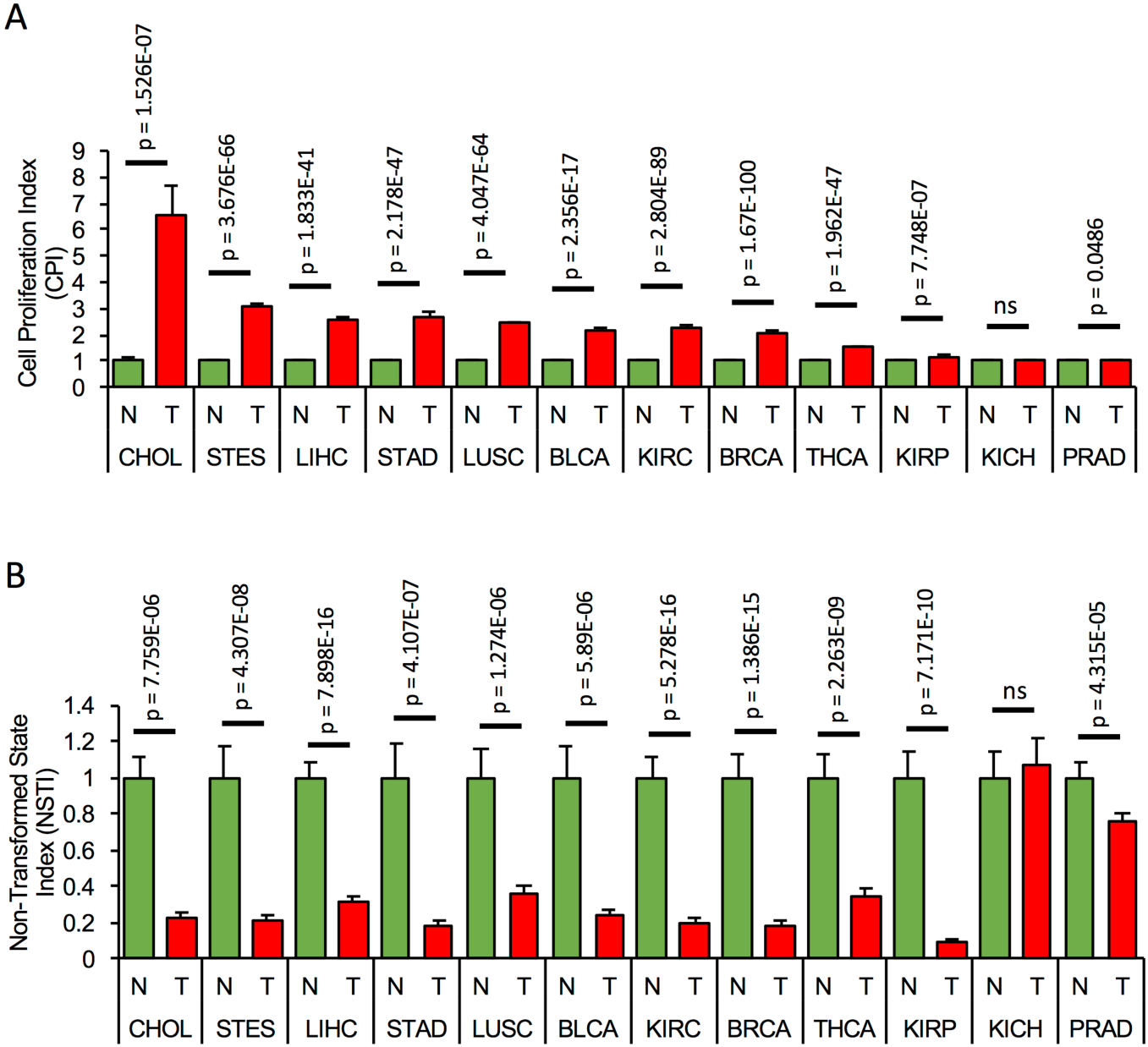
Concomitant activation of the proliferation and transformation networks is cancer type-specific. (A) Cell proliferation index, *CPI*, and (B) Non-transformed state index, *NSTI*, calculated from the measured mRNA expression levels of the network components in the non-tumor, normal (N) samples, (green bars) and tumor condition (T, red bars) of 12 tumor cohorts from TCGA. In each case, mRNA levels are relative to the non-tumor condition. A list of all components used in the analysis can be found in Supplementary Table 5.

We concluded that concomitant activation of the cell proliferation and transformation networks is cancer type-specific and predominantly occurs in gastrointestinal cancers.

### Patient-to-patient heterogeneity in the activation of the proliferation and transformation networks

Principal component analysis (PCA) based on the expression of all network components was performed in three cohorts from TCGA: (1) the cholangiocarcinoma cohort, which displays the highest *CPI* levels; (2) the hepatocellular carcinoma cohort, which shows high *CPI* and low *NSTI* values, and (3) the prostate adenocarcinoma cohort characterized by low *CPI* and high *NSTI*. This analysis revealed that non-tumor (blue dots) and tumor samples (red dots) cluster separately in cholangiocarcinoma (Fig. 6A), suggesting that the combined expression of the components of both networks can be used as a proxy to determine the tumorigenic state of a sample in this cohort.

**Figure 6.**
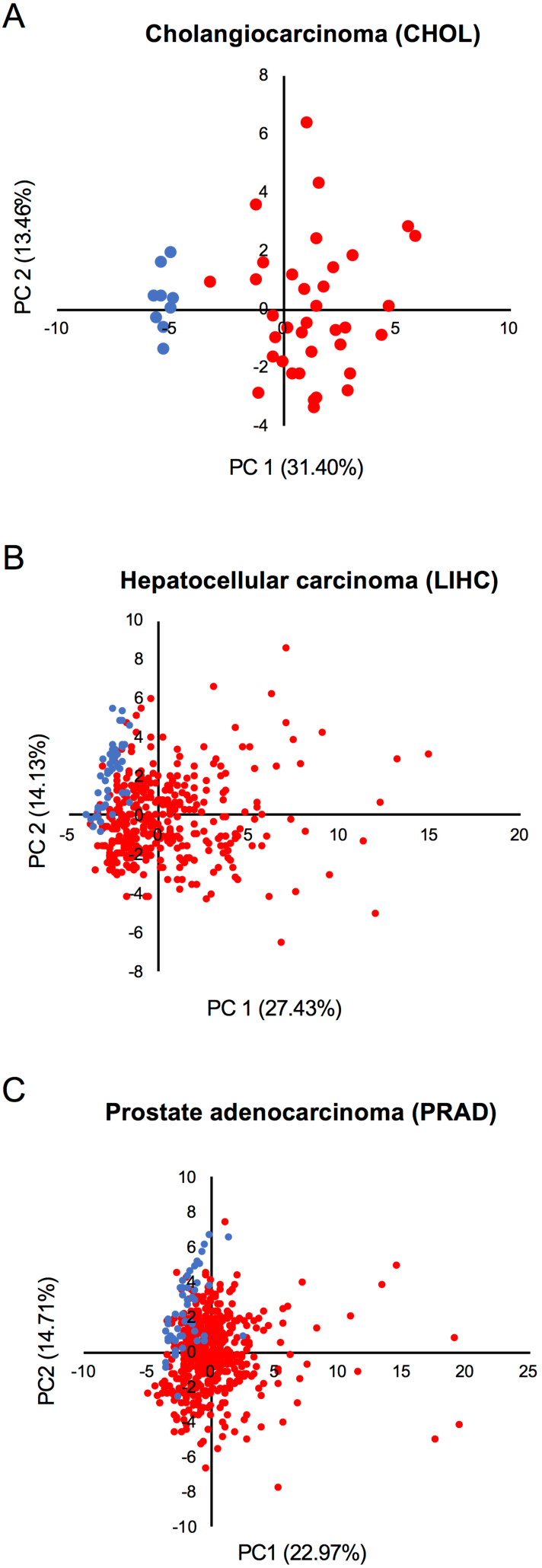
Activation of the networks in tumors is heterogeneous among patients. (A) PCA analysis based on the expression of all network components in the non-tumor (blue dots) and tumor condition (red dots) of (A) cholangiocarcinoma, (B) hepatocellular carcinoma, and (C) prostate adenocarcinoma cohort from TCGA.

A larger level of heterogeneity was detected within the hepatocellular carcinoma and prostate adenocarcinoma cohorts (Fig. 6B-C). Moreover, some tumors cluster together with normal samples indicating that the networks are not activated in all samples. Also, despite that the mean *CPI* and *NSTI* values of prostate adenocarcinoma do not significantly differ from those in corresponding non-tumor samples (Fig. 5A, B), some individual tumors are characterized by high activation of the networks (Fig. 6C).

We concluded that the network activation varies from patient to patient and depends on tumor-type.

### Dynamics of cell proliferation and cell transformation of a heterogeneous cell population

To determine the source of heterogeneity observed in patient samples, we incorporated a stochastic source of heterogeneity in a cell population model by applying, for each cell, uniform random variations around the basal value of each kinetic parameter. We plotted the simulated levels of *CPI* as a function of *NTSI* in a heterogeneous cell population of quiescent, non-transformed cells (Fig. 7A, C, E; orange dots correspond to *CPI* and *NTSI* in the absence of random variations on parameter’s values), and in a population of transformed, proliferative cells (Fig. 7B, D, F; green dots correspond to *CPI* and *NTSI* in the absence of random variations). 10%, 25% and 50% of uniform random variations on each parameter were considered.

**Figure 7.**
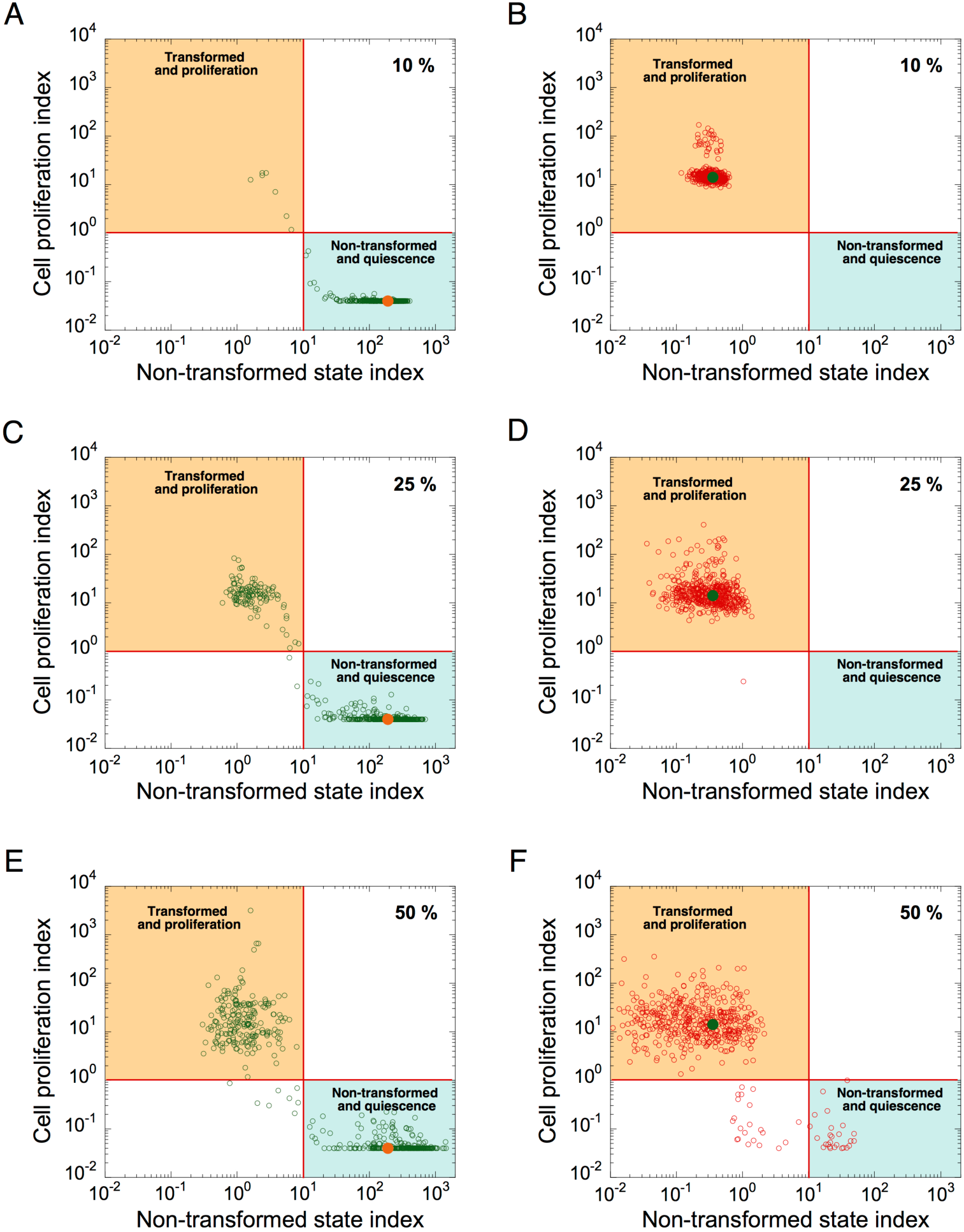
Modeling malignant cell transformation and cell proliferation distributions in a heterogeneous cell population. *CPI versus NSTI* is shown in a heterogeneous cell population starting from a non-transformed, quiescent, cell state (A, C, E) or from a transformed, proliferative, cell state (B, D, F). In both states, 10% (A, B), 25% (C, D), or 50% (E, F) of uniform random variations from the basal value of each parameter are considered. Each circle corresponds to one cell in a population of 500 cells. Horizontal lines define an arbitrary threshold value for *CPI*, which is equal to 1, above which cells are considered into proliferation; vertical lines define an arbitrary threshold value for *NSTI*, equal to 10, above which cells are considered in a non-transformed state. Orange (A, C, E) and green dots (B, D, F) correspond to the value of both indexes in the absence of random variation on parameters. Initial conditions are given in Supplementary Table 4. Basal parameter values are as in Supplementary Table 3.

Simulations indicate that the non-transformed, quiescent, cell state is less robust to random fluctuations than the transformed, proliferative, state (compare Figs. 7C with 7D, and 7E with 7F). Indeed, starting in a non-transformed, quiescent, cell state, a large proportion of cells switch to a transformed, proliferative, state in the presence of random fluctuations in gene expression (Fig. 7C and 7E). However, from a transformed, proliferative, cell state, only a small proportion of cells switch to a quiescent and non-transformed state (Fig. 7F).

Thus, random fluctuations in gene expression could trigger abrupt switches in the dynamics of the cell cycling and transformation networks. “Non-tumor” state (quiescent, non-transformed cells) is more sensitive to random fluctuations in gene expression than “tumor” state (transformed, proliferative state).

We next assessed if the levels of key network components, i.e. Let-7, PTEN, or Ras, may affect the robustness of these cell states. In a heterogeneous population of non-transformed, quiescent, cells with 25% of random parameter variations (Supplementary Figs. 4A-F), an increase in Let-7 strengthens the robustness of the non-transformed, quiescent, cell state towards random fluctuations in gene expression and prevents random switches to a transformed, proliferative, cell state (compare basal conditions in Supplementary Fig. 4A with Supplementary Figs. 4B, C, D where *V*_SLET7_ changes from 10 to 12, 20, and 50, respectively). Similarly, an increase in PTEN or a decrease in Ras also improved the robustness of the non-transformed, quiescent, cell state (compare Supplementary Figs. 4A with 4E and 4F).

Along the same lines, tumor suppressors and oncogenes can also impact on the robustness of the transformed, proliferative, cell state (Supplementary Fig. 4G-L). Some cells revert to a non-transformed, quiescent, state following a large increase in Let-7 (compare Supplementary Fig. 4G with 4H, I, J). However, an increase in PTEN or a decrease in Ras (similar to the conditions in Supplementary Figs. 4E and 4F) are unable to revert cells to a non-transformed, quiescent, state (compare Supplementary Fig. 4G with Figs. 4K and 4L).

Thus, an increase in tumor suppressors or a decrease in oncogenes reduces the probability of stochastic switches to a transformed, proliferative, cell state. However, if cells are already in a proliferative, transformed state, similar changes in tumor suppressors or oncogenes do not permit to revert back to a more “healthy” cell phenotype, which highlights an irreversible process in cancer progression.

Finally, since cholangiocarcinomas are characterized by strong network activation (Figs. 5 and 6A), we analyzed if the cell population model can qualitatively reproduce the networks’ switch from normal to tumor condition. Plotting cyclin B1 mRNA as a function of cyclin E1 or Let-7c RNA (Fig. 8A-D), and plotting Kras mRNA (representative as Ras) as a function of Let-7c (Fig. 8E, F) revealed expression profiles that are qualitatively very similar to those predicted by the mathematical model of a heterogeneous cell population (Fig. 8B, D, E). We concluded that the cell population model can be used to assess the qualitative dynamics of the switch of both networks in cholangiocarcinomas.

**Figure 8.**
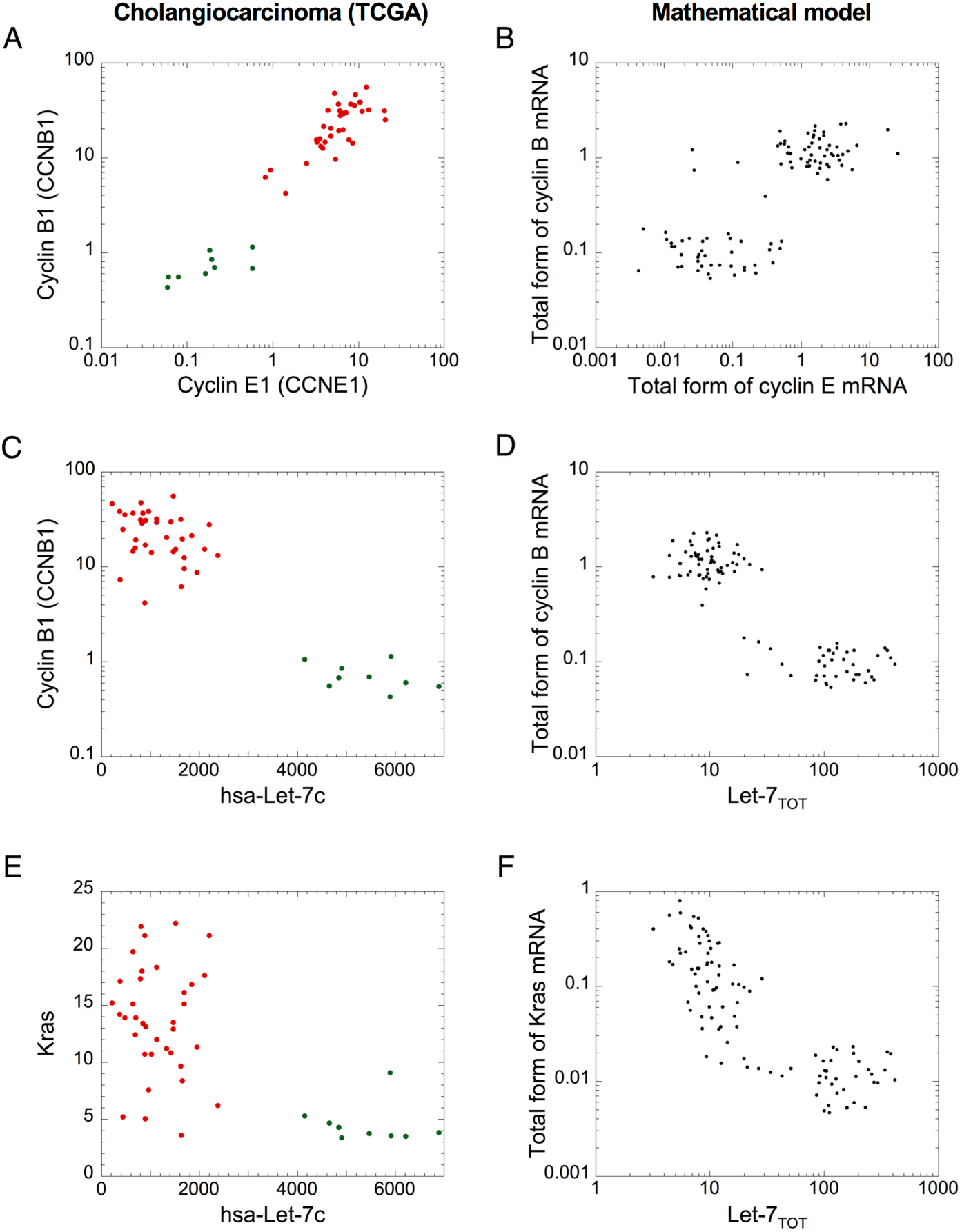
A cell population model assesses the qualitative dynamics of the network switches in cholangiocarcinoma. mRNA levels of (A, B) cyclin B1 *versus* cyclin E1, (C, D) cyclin B1 *versus* Let-7c, and (E, F) Kras *versus* Let-7c of cholangiocarcinoma cohort from TCGA (A, C, E), and (B, D, F) in a model for a heterogeneous cell population where 50% of uniform random variations are considered around the basal value of each parameter. (B, D, F) Simulations are performed with a population of 100 cells. Initial conditions are given in Supplementary Table 4. Src = 0.0000001 and other parameter values are as in Supplementary Table 3. (A, C, E) green dots: non-tumor samples (n = 9), red dots: tumor samples (n = 36).

## Discussion

Tumorigenesis rests on many biological features which include sustained proliferative signaling, evading growth suppressors, resisting cell death, promoting angiogenesis, ensuring replicative immortality and eliciting invasion and metastasis (Hanahan and Weinberg, 2011). Here, we built a mathematical model to analyze the dynamical properties of a Let-7-dependent mechanism coupling cell proliferation and an epigenetic switch driving malignant transformation.

Our mathematical model illustrates qualitatively how Cdk-dependent and transformation networks may interact, and proposes a mechanism, acting through Let-7, which suggests that cyclin overexpression can promote cell proliferation, while inducing and accelerating malignant transformation. Indeed, overexpression of cyclins progressively sponges the free form of Let-7. The latter will no longer be available to repress the components of the transformation network, leading to the activation of the epigenetic switch. This effect is known as competing-endogenous, ceRNA, effect. CeRNAs regulates other RNA transcripts by competing for shared miRNA and were involved in tumorigenesis (Salmena et al., 2011;Tay et al., 2014;Chiu et al., 2017). Here Let-7 is the shared miRNA between both networks. Let-7 was shown to be involved in different ceRNA mechanisms. Indeed, Let-7e can modulate the inflammatory response in vascular endothelial cells through a ceRNA effect (Lin et al., 2017). Imprinted H19 lncRNA, which plays important roles in development, cancer and metabolism, modulates Let-7 availability by acting as a molecular sponge and causing precocious muscle differentiation (Kallen et al., 2013). Moreover, amplification of MYCN mRNA levels in neuroblastoma can sponge Let-7 thereby rendering LIN28B dispensable for cancer progression (Powers et al., 2016). Note however that experimental and theoretical studies indicate that a ceRNA effect between multiple RNA transcripts and the shared miRNA is effective only in the presence of adequate expression levels of the transcripts and the miRNA (Gerard and Novak, 2013;Denzler et al., 2014;Tay et al., 2014).

Our mathematical model further shows that random variations in gene expression from cell to cell can create large fluctuations in the global network dynamics leading to stochastic switches of some cells to a transformed and proliferative state. These stochastic switches in the GRN dynamics could be a source of heterogeneity in cancer cell populations (Tang, 2012;Patel et al., 2014). Stochastic switches in gene networks were identified in hematopoietic tumor stem cells (Dingli et al., 2007), in the appearance of mammary tumor in mice (Bouchard et al., 1989), and in the differentiation and maturation of T lymphocytes (Davis et al., 1993). These switches may also give rise to an equilibrium in population of cancer cells (Gupta et al., 2011). Transcriptomic analysis of TCGA data suggests that the coupling between the cell cycle and a malignant transformation networks, and the activation of these networks in tumors are cancer type-specific, with predominant activation in gastrointestinal cancers. Our PCA analysis reveals inter-patient heterogeneity in network activation in tumors, which stresses the need to consider patient-specific characteristics when optimizing therapeutic strategies by reversing network dynamics of activated GRNs (Biankin et al., 2015).

In conclusion, by means of transcriptomic data analysis and modeling-based investigations, we identified a Let-7-dependent connection between two major GRNs involved in tumorigenesis, and whose activation is cancer-and patient-specific. We anticipate that a better characterization of the dynamics resulting from the combination of other GRNs specific for each patient will help providing a global GRN activation map for personalizing and optimizing cancer treatment.

## Acknowledgments

This work was supported by the “Fonds National de la Recherche Scientifique” (FNRS, Belgium) [Convention no T.0015.16 (26021674)].

## Supplementary Information

### Description of the mathematical model

The model is described by a set of 15 kinetic equations for the Cdk network driving the mammalian cell cycle and 14 kinetics equations for the inflammatory circuit that controls the dynamics of malignant transformation. Each equation represents the temporal evolution of the expression level of one component of the network. The different variables of the model are defined in Table S1, the kinetics equations are found in Table S2, while the description of the parameters and their numerical values used in the simulations are in Table S3.

In the model, we consider for simplicity a constant total concentration of NF-κB (*NFKB*_TOT_), E2F (E2F_TOT_), and APC (APC_TOT_). Activation and inactivation reactions of E2F in Eq. [5], APC in Eq. [15] and NF-κB in Eq. [16] behave as Goldbeter-Koshland switches (Goldbeter and Koshland, 1981). All other processes of the model rest on mass-action kinetics. Moreover, to couple the model for the Cdk network with the model for the inflammatory circuit leading to malignant transformation, equations [1] and [18] have been replaced by equation [18’].

### Numerical simulations and PCA analysis

Numerical simulations were performed with XPPAUTO (http://www.math.pitt.edu/~bard/xpp/xpp.html) and matlab. PCA analysis were done with FactoMineR (Le, 2008).

### Analysis of RNASeq data

RNASeq and miRNASeq data of the different patient cohorts were from TCGA database (http://firebrowse.org/). For each cohort, we converted the “scaled_estimate” in the “illuminahiseq_rnaseqv2_unc_edu_Level_3_RSEM_genes” file into TPM by multiplying by 10^6^.

**Table S1.** Variables of the mathematical model

| <u>Name</u> | <u>Definition</u> |
| --- | --- |
| ***Model for the Cdk network, regulated by Let-7, driving the mammalian cell cycle*** |  |
| Let7 | MicroRNA family Let-7 |
| mMd | Cyclin D messenger RNA |
| Md | Active form of cyclin D/Cdk-4-6 complex |
| mMdlet7 | Inactive complex between cyclin D messenger RNA (mMd) and Let7 |
| E2F | Active form of the transcription factor E2F |
| mMe | Cyclin E messenger RNA |
| Me | Active form of cyclin E/Cdk2 complex |
| mMelet7 | Inactive complex between cyclin E messenger RNA (mMe) and Let7 |
| mMa | Cyclin A messenger RNA |
| Ma | Active form of cyclin A/Cdk2 complex |
| mMalet7 | Inactive complex between cyclin A messenger RNA (mMa) and Let7 |
| mMb | Cyclin B messenger RNA |
| Mb | Active form of cyclin B/Cdk1 complex |
| mMblet7 | Inactive complex between cyclin B messenger RNA (mba) and Let7 |
| APC | Anaphase-promoting complex regulating cyclin degradation |
| ***Model for the inflammatory circuit driving the epigenetic switch to cell transformation*** |  |
| Let7 | MicroRNA family Let-7 |
| NFKB | Transcription factor NF-κB |
| Lin28 | RNA binding protein Lin28 |
| mIL6 | IL6 messenger RNA |
| mIL6Let7 | Complex form between mIL6 and Let7 |
| IL6 | Interleukin 6 protein, which promotes the epigenetic switch leading to cell transformation |
| mRas | Ras oncogene messenger RNA |
| mRasLet7 | Complex form between mRas and Let7 |
| Ras | Protein form of Ras oncogene |
| STAT3 | Transcription factor STAT3, whose activity is crucial to promote cell transformation |
| miR21 | MicroRNA miR21, which represses the translation of the tumor suppressor PTEN |
| mPTEN | PTEN messenger RNA |
| miRmpten | Complex form between miR21 and mPTEN |
| PTEN | Protein form of the tumor suppressor PTEN (phosphatase and tensin homologue) |

**Table S2.**
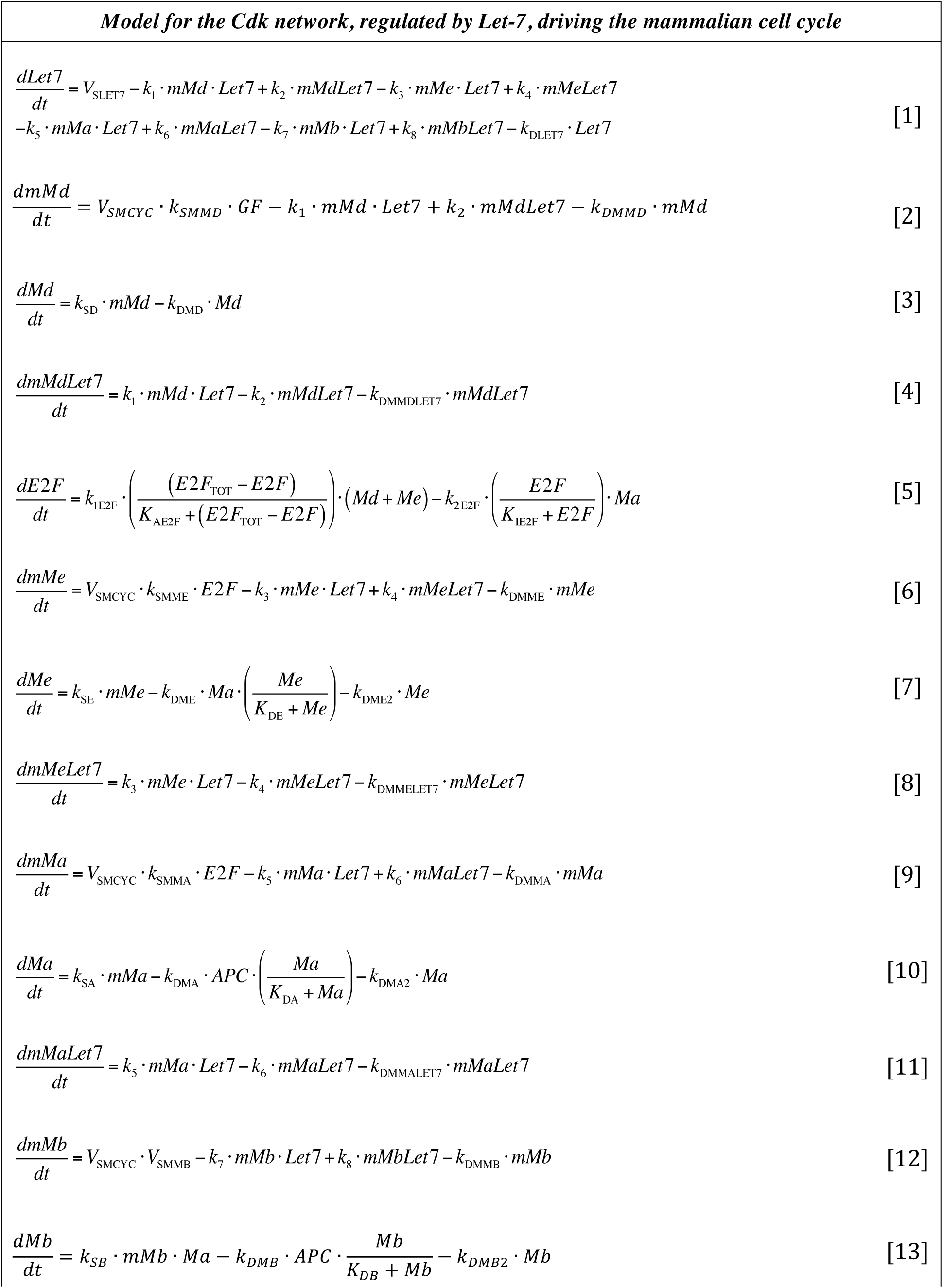
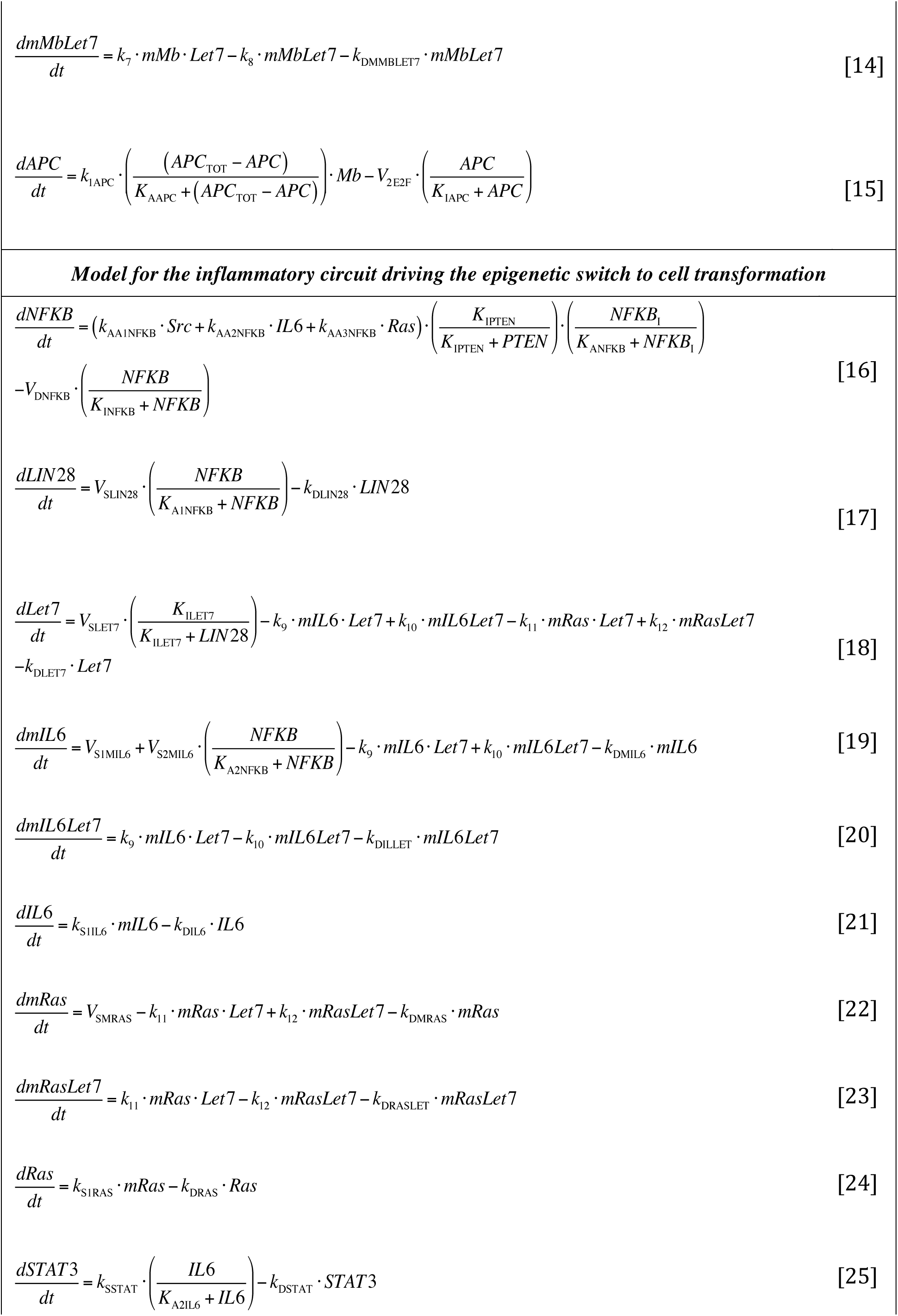
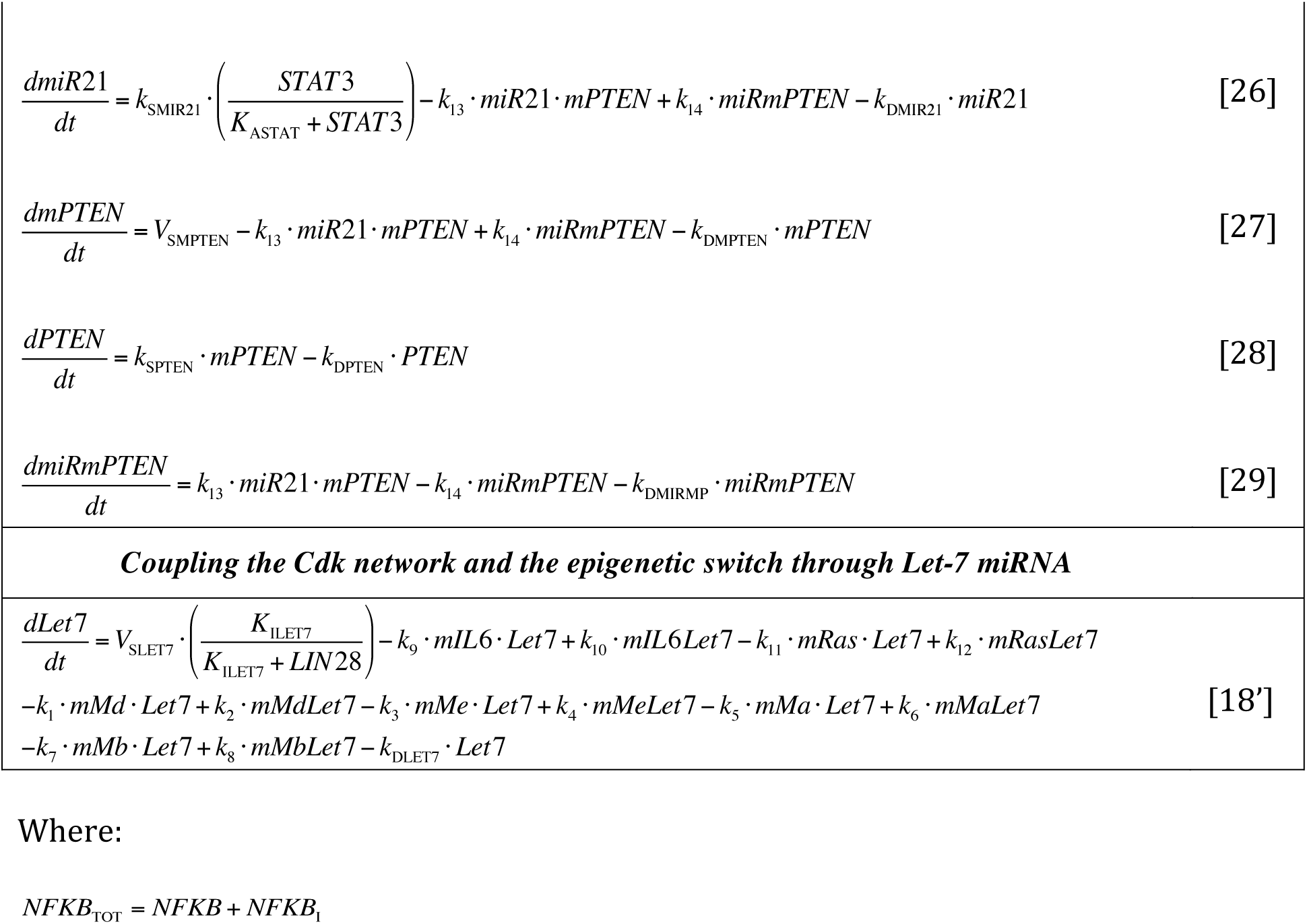
Kinetic equations of the mathematical model

**Table S3.**
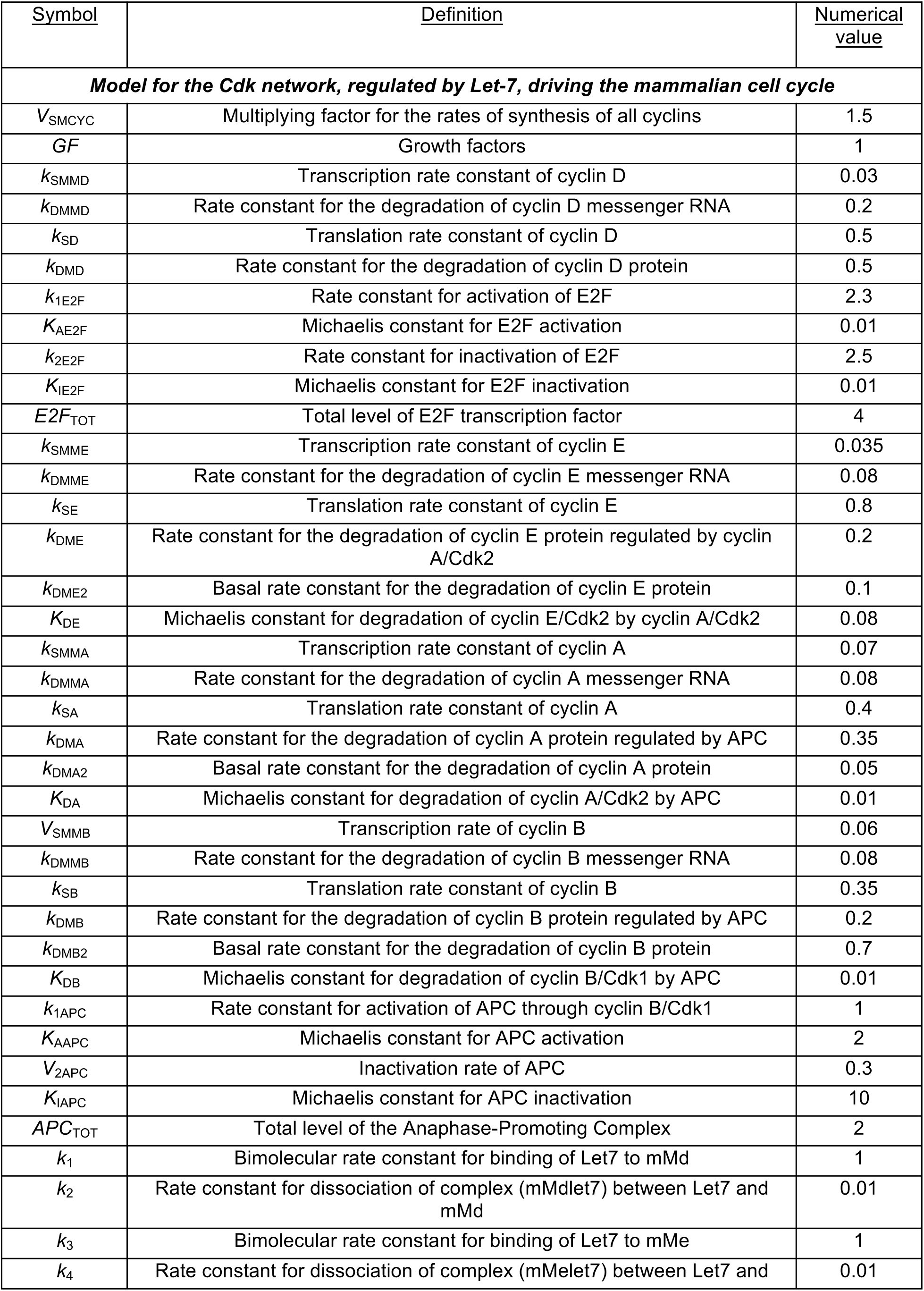
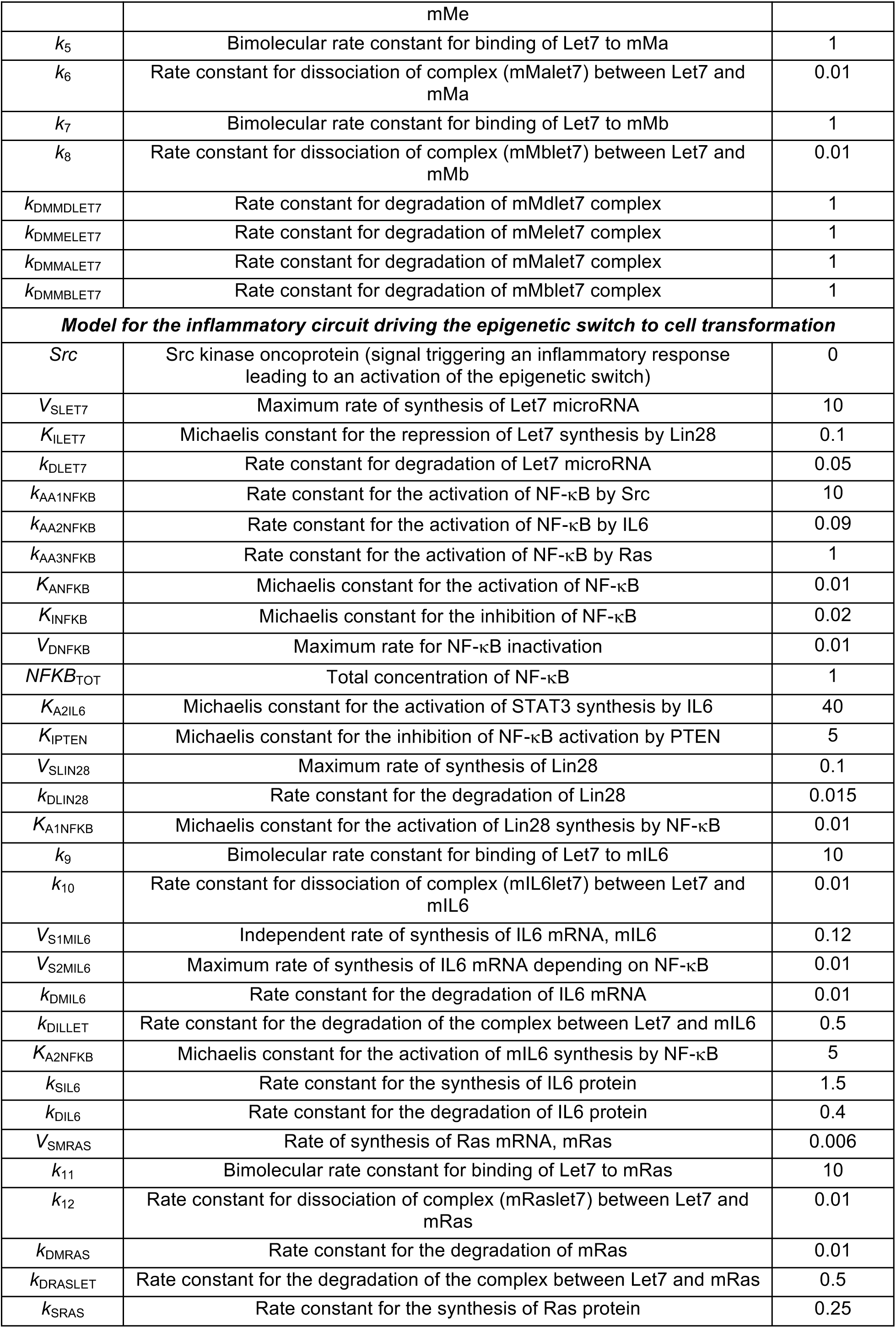
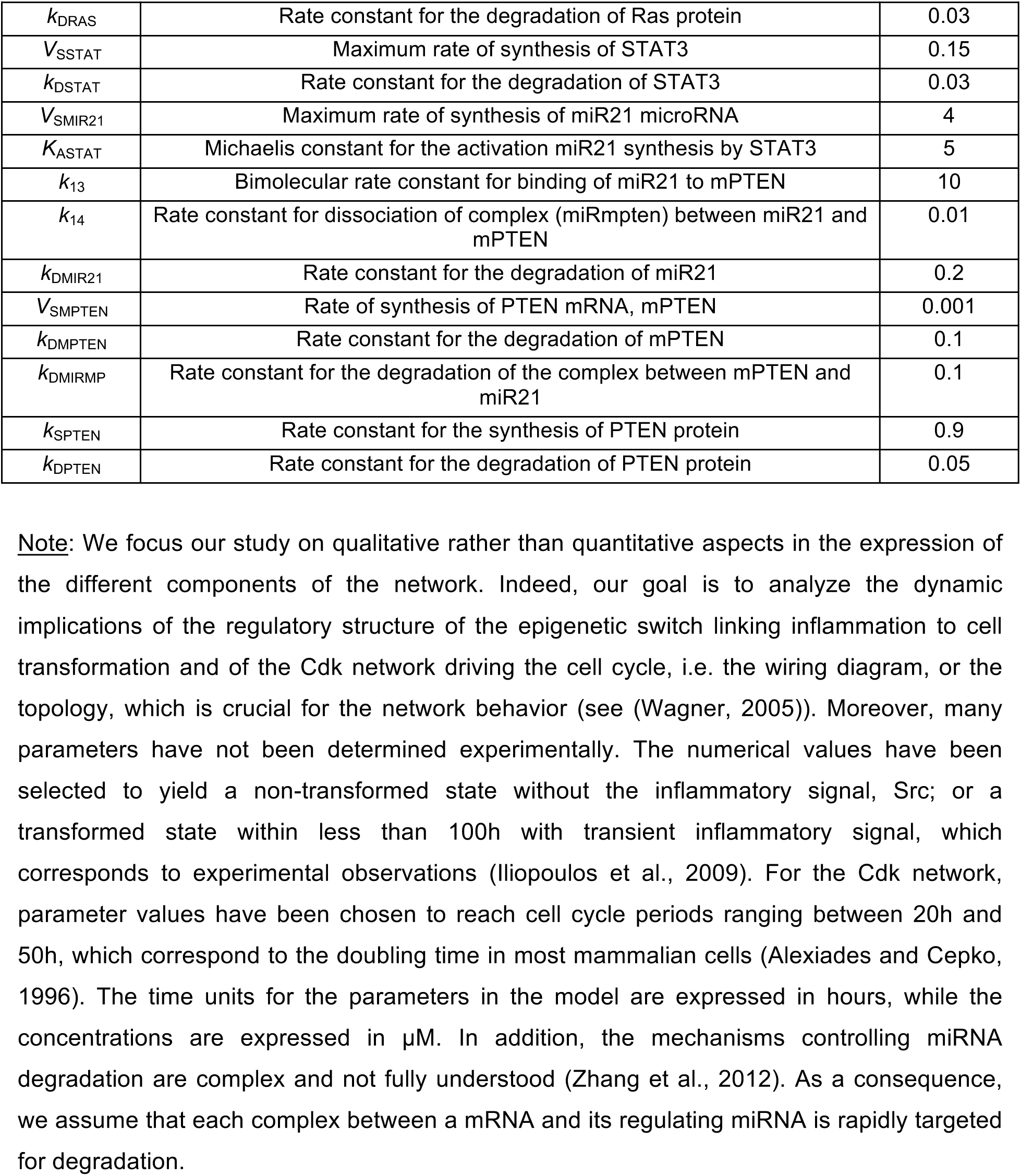
Parameters of the model

**Table S4.** Initial conditions

| **Variable** | **C.I. # 1**<br>**(leads to non-**<br>**transformed cell)** | **C.I. # 2**<br>**(leads to**<br>**transformed cell)** | **C.I. # 3**<br>**(cell population,**<br>**Figs. 8B,D,F)** |
| --- | --- | --- | --- |
| *NFKB* | 0.00045 | 0.5 | 0.01 |
| *Lin28* | 0.34 | 0.5 | 0.01 |
| *Let7* | 40 | 1 | 0 |
| *mIL6* | 0.0003 | 0.5 | 0.01 |
| *IL6* | 0.001 | 0.5 | 0.01 |
| *mRas* | 0.00001 | 0.5 | 0.01 |
| *Ras* | 0.0001 | 0.5 | 0.01 |
| *STAT3* | 0.0001 | 0.5 | 0.01 |
| *mPTEN* | 0.01 | 0 | 0.01 |
| *mRasLet7* | 0 | 0 | 0 |
| *mIL6Let7* | 0 | 0 | 0 |
| *miR21* | 0.0003 | 0 | 0 |
| *PTEN* | 0.17 | 0 | 0 |
| *MiRmPTEN* | 0 | 0 | 0 |

**Table S5.** List of the components used in the human tumors analysis (Figs. 5-6)

| # of components | Symbol | Name |
| --- | --- | --- |
| 1 | CCNA1 | Cyclin A1 |
| 2 | CCNB1 | Cyclin B1 |
| 3 | CCND1 | Cyclin A1 |
| 4 | CCNE1 | Cyclin E1 |
| 5 | CDK1 | Cyclin-dependent kinase 1 |
| 6 | CDK2 | Cyclin-dependent kinase 2 |
| 7 | CDK4 | Cyclin-dependent kinase 4 |
| 8 | CDK6 | Cyclin-dependent kinase 6 |
| 9 | E2F1 | E2F transcription factor 1 |
| 10 | E2F2 | E2F transcription factor 2 |
| 11 | E2F3 | E2F transcription factor 3 |
| 12 | E2F4 | E2F transcription factor 4 |
| 13 | E2F5 | E2F transcription factor 5 |
| 14 | E2F6 | E2F transcription factor 6 |
| 15 | E2F7 | E2F transcription factor 7 |
| 16 | E2F8 | E2F transcription factor 8 |
| 17 | IL6R | Interleukin 6 receptor |
| 18 | IL6 | Interleukin 6 |
| 19 | KRAS | KRAS oncogene |
| 20 | LIN28A | Lin-28 homolog A |
| 21 | LIN28B | Lin-28 homolog B |
| 22 | PTEN | Phosphatase and tensin homolog |
| 23 | STAT3 | Signal transducer and activator of transcription 3 |
| 24 | hsa-let-7a-1 | MicroRNA let-7a-1 |
| 25 | hsa-let-7a-2 | MicroRNA let-7a-2 |
| 26 | hsa-let-7a-3 | MicroRNA let-7a-3 |
| 27 | hsa-let-7b | MicroRNA let-7b |
| 28 | hsa-let-7c | MicroRNA let-7c |
| 29 | hsa-let-7d | MicroRNA let-7d |
| 30 | hsa-let-7e | MicroRNA let-7e |
| 31 | hsa-mir-21 | MicroRNA 21 |

**Figure S1.**
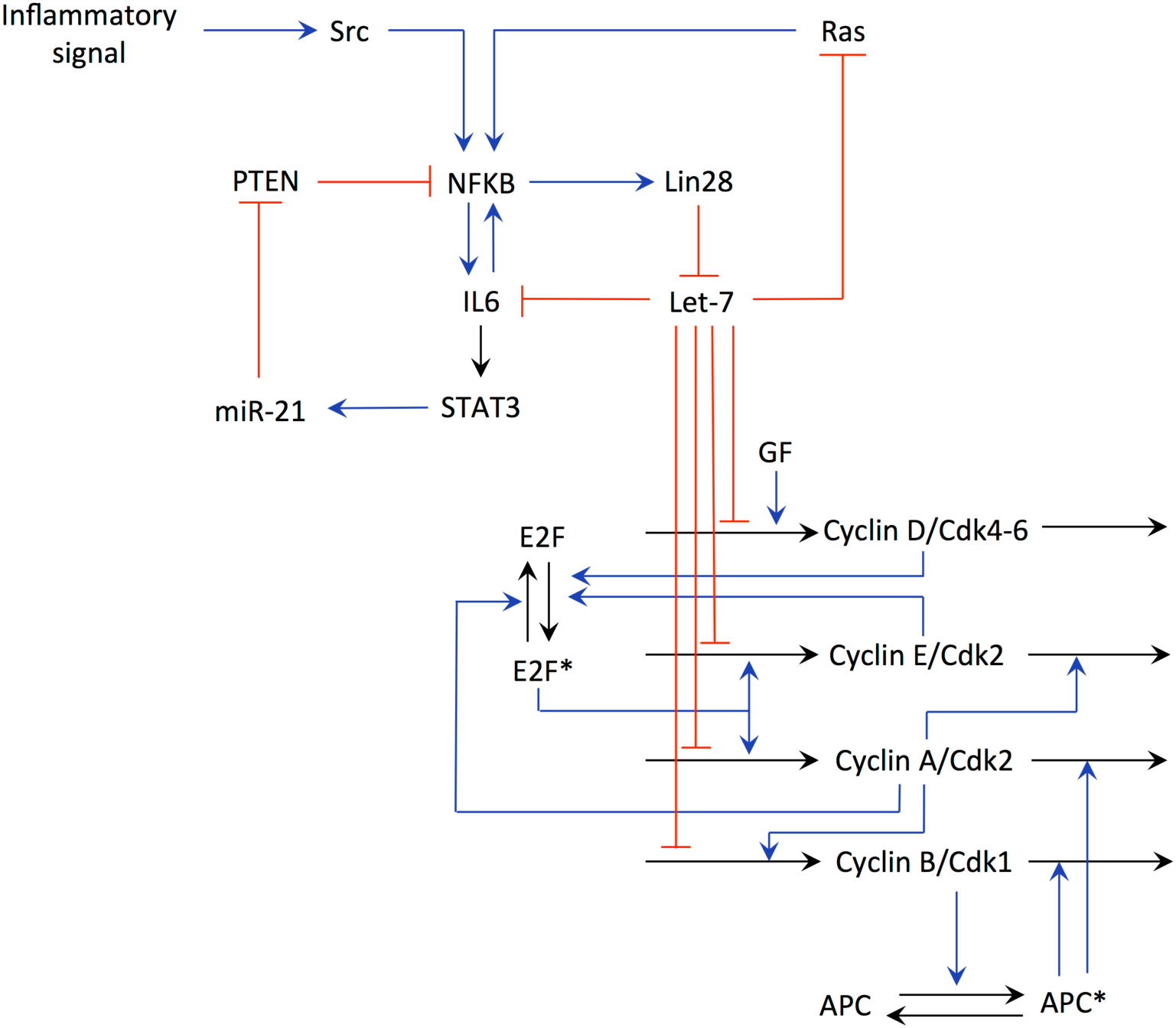
Detailed scheme of the model. The model for the epigenetic switch linking inflammation to cell transformation originates from a previous study (Gerard et al., 2014), while the model for the post-transcriptional regulation of the Cdk network by Let-7 is based on a skeleton model for the Cdk network driving the mammalian cell cycle (Gerard and Goldbeter, 2011). Let-7, which is involved in multiple positive feedback loops, can repress the synthesis of each cyclin.

**Figure S2.**
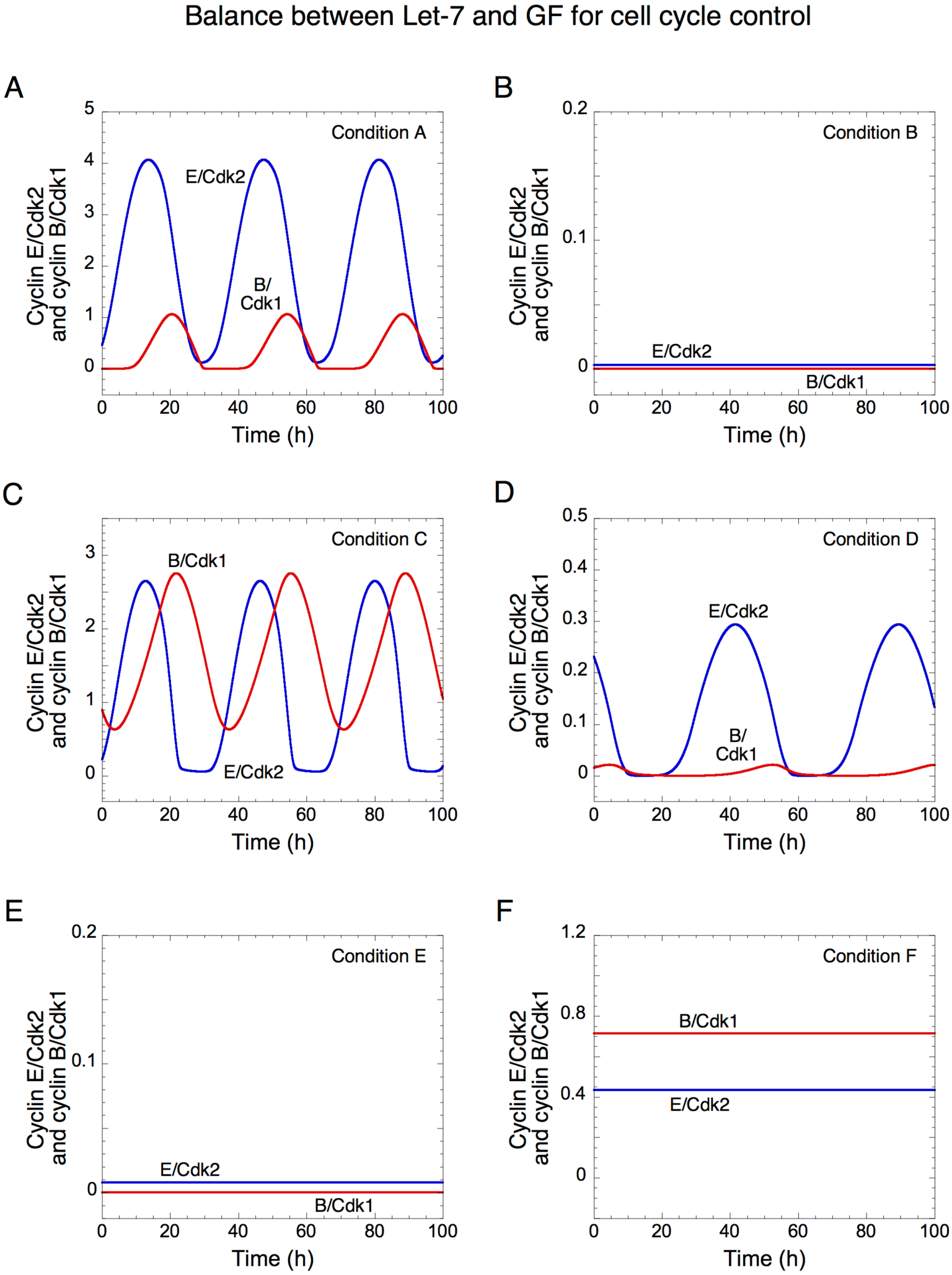
Cell proliferation versus cell cycle arrest mediated by Let-7 and GF levels. The temporal evolutions of cyclin E/Cdk2 (blue) and cyclin B/Cdk1 (red) in panels A to F correspond to conditions A to F of panel A from Fig. 2. In A, *GF* = 3 and *V*_SLET7_ = 0.25; in B, *GF* = 3 and *V*_SLET7_ = 2; in C, *GF* = 25 and *V*_SLET7_ = 0.25; in D, *GF* = 25 and *V*_SLET7_ = 1.6; in E, *GF* = 25 and *V*_SLET7_ = 5; and in F, *GF* = 70 and *V*_SLET7_ = 1.5. Other parameter values are as in Table S3.

**Figure S3.**
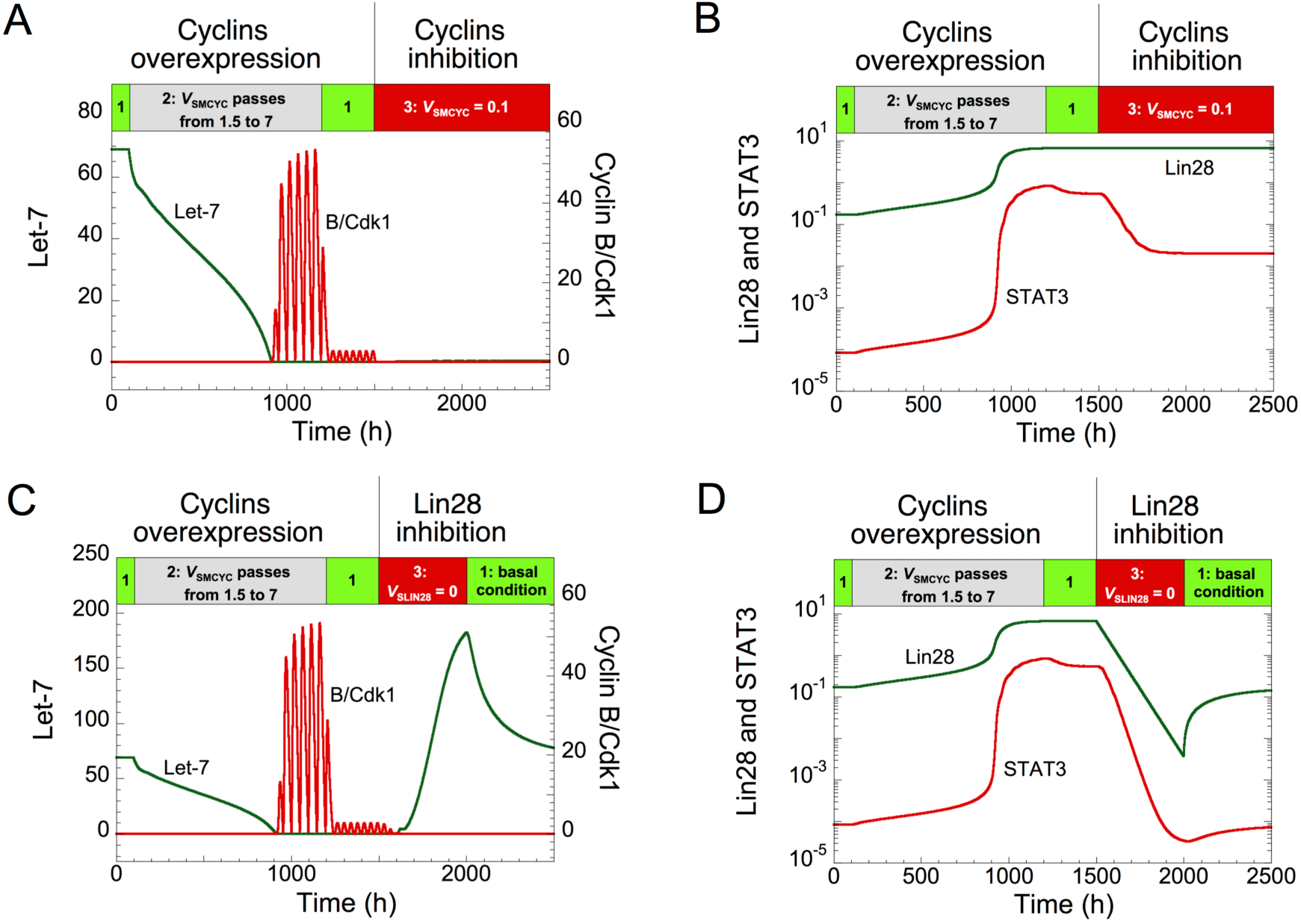
Effect of the Cdk network on the dynamics of the transformation network. (A, C) Temporal evolution of Let-7 and cyclin B/Cdk1. From a non-transformed and quiescent cell state, *V*_SMCYC_ changes from 1.5 to 7 (at t > 100h), and promotes the epigenetic switch and cell proliferation. For 1200 h < t < 1500 h, *V*_SMCYC_ is set back to its initial value (= 1.5), cell proliferation and the transformed cell state are maintained, which is characterized by sustained oscillations in cyclin B/Cdk1 and low levels in Let-7. In A, at t = 1500h, *V*_SMCYC_ changes from 1.5 to 0.1, preventing cell proliferation without recovering a non-transformed cell state since cyclin B/Cdk1 tends to a low stable steady state level, while Let-7 levels remain low. In C, for 1500h < t < 2000h, the synthesis rate of Lin28, *V*_SLIN28_ is set to 0, allowing the recovery of a stable non-transformed state defined by high levels of Let-7 and impeding cell proliferation, characterized by a low, stable steady state, level of cyclin B/Cdk1. In C, for t > 2000h, basal conditions are used. (B, D) Temporal evolution of Lin28 and STAT3 corresponds to conditions of panels A and C, respectively. Other parameter values are as in Table S3, which correspond to basal conditions.

**Figure S4.**
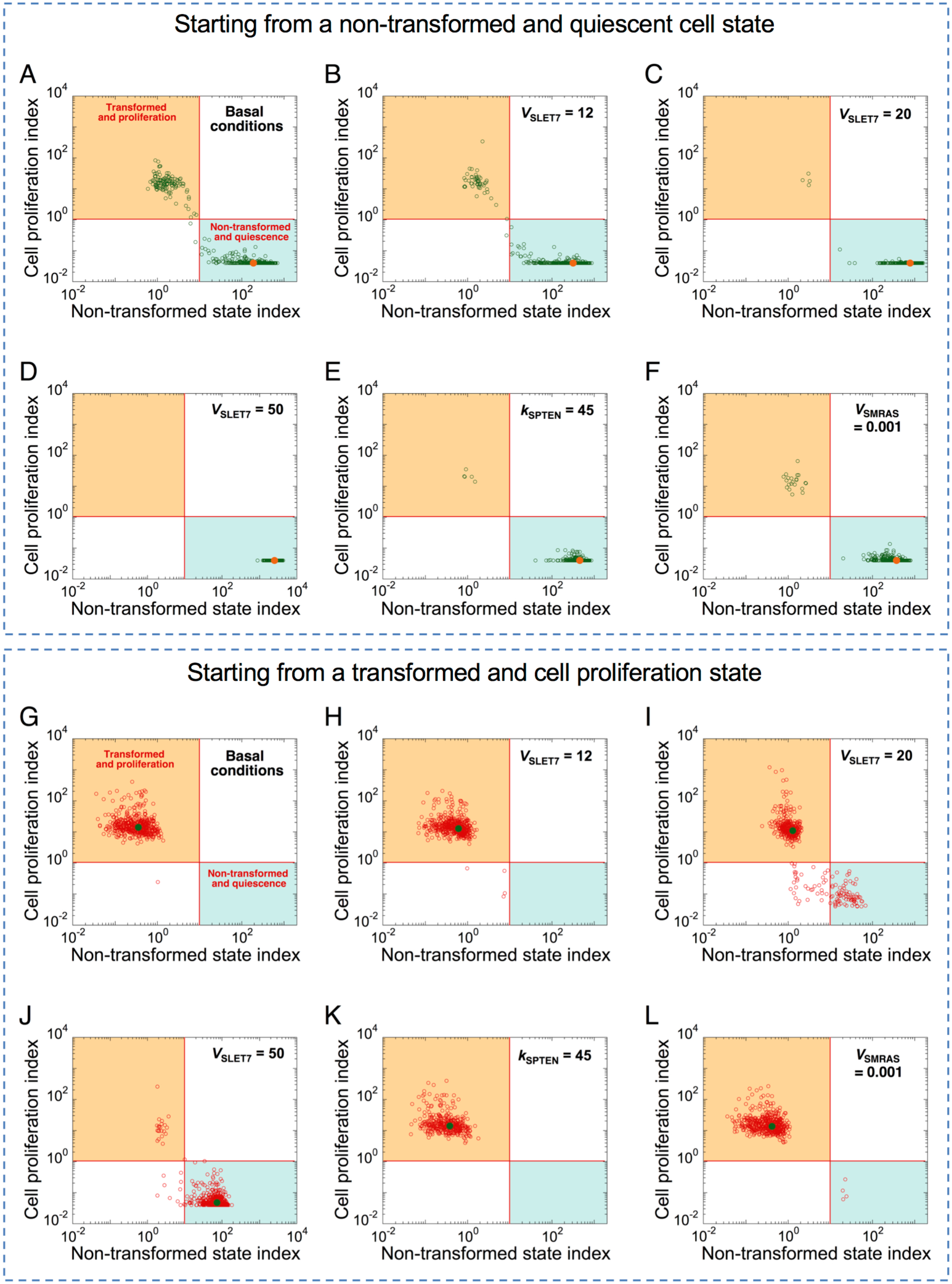
Effect of oncogenes and tumor suppressors on the robustness of the dynamics. Starting from a non-transformed and quiescent cell state (A-F), or from a transformed and proliferating cell state (G-L), the cell proliferation index *versus* non-transformed state index is shown in a heterogeneous cell population with 25% of uniformed random variation of all parameters of the model. In both cases, simulations are performed in the basal conditions (A and G); in the presence of various overexpression levels of Let-7: *V*_SLET7_ = 12 in B and H, 20 in C and I, and 50 in D and J; in the presence of PTEN overexpression, *k*_SPTEN_ = 45 in E and K, or in the presence of Ras inhibition, *V*_SMRAS_ = 0.001 instead of 0.006 in F and L. In each case, circles correspond to one cell in a population of 500 cells. The orange (A-F) and green dots (G-L) correspond to the levels of *CPI* and *NTSI* in the absence of random variation on parameters. Basal condition corresponds to the parameter values in Table S3.

## References

Biankin, A.V., Piantadosi, S., and Hollingsworth, S.J. (2015). Patient-centric trials for therapeutic development in precision oncology. Nature 526, 361–370.

Bouchard, L., Lamarre, L., Tremblay, P.J., and Jolicoeur, P. (1989). Stochastic appearance of mammary tumors in transgenic mice carrying the MMTV/c-neu oncogene. Cell 57, 931–936.

Bueno, M.J., and Malumbres, M. (2011). MicroRNAs and the cell cycle. Biochim Biophys Acta 1812, 592–601.

Chiu, H.S., Martinez, M.R., Bansal, M., Subramanian, A., Golub, T.R., Yang, X., Sumazin, P., and Califano, A. (2017). High-throughput validation of ceRNA regulatory networks. BMC Genomics 18, 418.

Davis, C.B., Killeen, N., Crooks, M.E., Raulet, D., and Littman, D.R. (1993). Evidence for a stochastic mechanism in the differentiation of mature subsets of T lymphocytes. Cell 73, 237–247.

Denzler, R., Agarwal, V., Stefano, J., Bartel, D.P., and Stoffel, M. (2014). Assessing the ceRNA hypothesis with quantitative measurements of miRNA and target abundance. Mol Cell 54, 766–776.

Dingli, D., Traulsen, A., and Pacheco, J.M. (2007). Stochastic dynamics of hematopoietic tumor stem cells. Cell Cycle 6, 461–466.

Edgar, B.A., Zielke, N., and Gutierrez, C. (2014). Endocycles: a recurrent evolutionary innovation for post-mitotic cell growth. Nat Rev Mol Cell Biol 15, 197–210.

Fofaria, N.M., and Srivastava, S.K. (2015). STAT3 induces anoikis resistance, promotes cell invasion and metastatic potential in pancreatic cancer cells. Carcinogenesis 36, 142–150.

Gerard, C., and Goldbeter, A. (2009). Temporal self-organization of the cyclin/Cdk network driving the mammalian cell cycle. Proc Natl Acad Sci U S A 106, 21643–21648.

Gerard, C., and Goldbeter, A. (2011). A skeleton model for the network of cyclin-dependent kinases driving the mammalian cell cycle. Interface Focus 1, 24–35.

Gerard, C., Gonze, D., Lemaigre, F., and Novak, B. (2014). A model for the epigenetic switch linking inflammation to cell transformation: deterministic and stochastic approaches. PLoS Comput Biol 10, e1003455.

Gerard, C., and Novak, B. (2013). microRNA as a potential vector for the propagation of robustness in protein expression and oscillatory dynamics within a ceRNA network. PLoS One 8, e83372.

Gillett, C., Fantl, V., Smith, R., Fisher, C., Bartek, J., Dickson, C., Barnes, D., and Peters, G. (1994). Amplification and overexpression of cyclin D1 in breast cancer detected by immunohistochemical staining. Cancer Res 54, 1812–1817.

Gupta, P.B., Fillmore, C.M., Jiang, G., Shapira, S.D., Tao, K., Kuperwasser, C., and Lander, E.S. (2011). Stochastic state transitions give rise to phenotypic equilibrium in populations of cancer cells. Cell 146, 633–644.

Hanahan, D., and Weinberg, R.A. (2011). Hallmarks of cancer: the next generation. Cell 144, 646–674.

Iliopoulos, D., Hirsch, H.A., and Struhl, K. (2009). An epigenetic switch involving NF-kappaB, Lin28, Let-7 MicroRNA, and IL6 links inflammation to cell transformation. Cell 139, 693–706.

Iliopoulos, D., Jaeger, S.A., Hirsch, H.A., Bulyk, M.L., and Struhl, K. (2010). STAT3 activation of miR-21 and miR-181b-1 via PTEN and CYLD are part of the epigenetic switch linking inflammation to cancer. Mol Cell 39, 493–506.

Johnson, C.D., Esquela-Kerscher, A., Stefani, G., Byrom, M., Kelnar, K., Ovcharenko, D., Wilson, M., Wang, X., Shelton, J., Shingara, J., Chin, L., Brown, D., and Slack, F.J. (2007). The let-7 microRNA represses cell proliferation pathways in human cells. Cancer Res 67, 7713–7722.

Kallen, A.N., Zhou, X.B., Xu, J., Qiao, C., Ma, J., Yan, L., Lu, L., Liu, C., Yi, J.S., Zhang, H., Min, W., Bennett, A.M., Gregory, R.I., Ding, Y., and Huang, Y. (2013). The imprinted H19 lncRNA antagonizes let-7 microRNAs. Mol Cell 52, 101–112.

Lin, Z., Ge, J., Wang, Z., Ren, J., Wang, X., Xiong, H., Gao, J., Zhang, Y., and Zhang, Q. (2017). Let-7e modulates the inflammatory response in vascular endothelial cells through ceRNA crosstalk. Sci Rep 7, 42498.

Morgan, D.O. (2007). The cell cycle: principles of control. London: New Science Press Ltd in association with Oxford University Press.

Novak, B., and Tyson, J.J. (2004). A model for restriction point control of the mammalian cell cycle. J Theor Biol 230, 563–579.

Patel, A.P., Tirosh, I., Trombetta, J.J., Shalek, A.K., Gillespie, S.M., Wakimoto, H., Cahill, D.P., Nahed, B.V., Curry, W.T., Martuza, R.L., Louis, D.N., Rozenblatt-Rosen, O., Suva, M.L., Regev, A., and Bernstein, B.E. (2014). Single-cell RNA-seq highlights intratumoral heterogeneity in primary glioblastoma. Science 344, 1396–1401.

Pok, S., Wen, V., Shackel, N., Alsop, A., Pyakurel, P., Fahrer, A., Farrell, G.C., and Teoh, N.C. (2013). Cyclin E facilitates dysplastic hepatocytes to bypass G1/S checkpoint in hepatocarcinogenesis. J Gastroenterol Hepatol 28, 1545–1554.

Powers, J.T., Tsanov, K.M., Pearson, D.S., Roels, F., Spina, C.S., Ebright, R., Seligson, M., De Soysa, Y., Cahan, P., Theissen, J., Tu, H.C., Han, A., Kurek, K.C., Lapier, G.S., Osborne, J.K., Ross, S.J., Cesana, M., Collins, J.J., Berthold, F., and Daley, G.Q. (2016). Multiple mechanisms disrupt the let-7 microRNA family in neuroblastoma. Nature 535, 246–251.

Salmena, L., Poliseno, L., Tay, Y., Kats, L., and Pandolfi, P.P. (2011). A ceRNA hypothesis: the Rosetta Stone of a hidden RNA language? Cell 146, 353–358.

Sotiropoulou, G., Pampalakis, G., Lianidou, E., and Mourelatos, Z. (2009). Emerging roles of microRNAs as molecular switches in the integrated circuit of the cancer cell. RNA 15, 1443–1461.

Tang, D.G. (2012). Understanding cancer stem cell heterogeneity and plasticity. Cell Res 22, 457–472.

Tay, Y., Rinn, J., and Pandolfi, P.P. (2014). The multilayered complexity of ceRNA crosstalk and competition. Nature 505, 344–352.

Zhu, X., Wu, L., Yao, J., Jiang, H., Wang, Q., Yang, Z., and Wu, F. (2015). MicroRNA let-7c Inhibits Cell Proliferation and Induces Cell Cycle Arrest by Targeting CDC25A in Human Hepatocellular Carcinoma. PLoS One 10, e0124266.

## Supplementary references

Alexiades, M.R., and Cepko, C. (1996). Quantitative analysis of proliferation and cell cycle length during development of the rat retina. Dev Dyn 205, 293–307.

Goldbeter, A., and Koshland, D.E., Jr. (1981). An amplified sensitivity arising from covalent modification in biological systems. Proc Natl Acad Sci U S A 78, 6840–6844.

Iliopoulos, D., Hirsch, H.A., and Struhl, K. (2009). An epigenetic switch involving NF-kappaB, Lin28, Let-7 MicroRNA, and IL6 links inflammation to cell transformation. Cell. 139, 693–706.

Le, S.J., J.; Husson, F. (2008). FactoMineR: An R Package for Multivariate Analysis. Journal of Statistical Software 25, 1–18.

Wagner, A. (2005). Circuit topology and the evolution of robustness in two-gene circadian oscillators. Proc Natl Acad Sci U S A 102, 11775–11780.

Zhang, Z., Qin, Y.W., Brewer, G., and Jing, Q. (2012). MicroRNA degradation and turnover: regulating the regulators. Wiley Interdiscip Rev RNA 3, 593–600.

